# Choice of phenotype scale is critical in biobank-based G×E tests

**DOI:** 10.64898/2026.01.20.694695

**Authors:** Manuela Costantino, Renée Fonseca, Zhengtong Liu, Zhenhong Huang, Sriram Sankararaman, Iain Mathieson, Andy Dahl

**Author notes:** Corresponding author: Manuela Costantino. These authors jointly supervised the work (Iain Mathieson, Andy Dahl).

## Abstract

The importance of gene-environment interactions (G×E) for complex human traits is heavily debated. Recently, biobank-based GWAS have revealed many statistically significant G×E signals, though most lack clear evidence of biological significance. Here, we partly explain this discrepancy by showing that many G×E signals simplify to additive effects on a different phenotype scale, a classical concern that is currently underappreciated. Our results clearly distinguish G×Sex effects on height, which vanish on the log scale, from G×Sex effects on testosterone, where the log scale uncovers biologically meaningful female-specific effects. Across 32 phenotypes in UK Biobank, we find that scaling by a power transformation can explain 46% of PGS×Sex interactions, and that simple log transformation can explain 23%, with similar results for other environments. We also show that phenotype scale can substantially impact GWAS discovery and the construction and evaluation of polygenic scores. Finally, we provide a set of guidelines to consider and choose phenotype scale in modern genetic studies.

## Introduction

Gene-environment (G×E) interaction models are a classical framework for studying how genetic architecture depends on the environment (1,2). G×E effects are well established in model systems, such as yeast fitness depending on growth medium or gene regulatory effects depending on chemical exposures (3–6). Examples of biomedically significant G×E have also been identified in complex human diseases, such as the sex-specific effect of *KLF14* variants on type 2 diabetes (7) and the 17q locus effect on asthma that depends on childhood rhinovirus illness (8).

Nonetheless, the importance of G×E for complex human traits generally remains unclear. Early attempts to identify G×E in candidate gene studies produced largely spurious results, but this can be simply attributed to underpowered studies (9,10). With the availability of biobank-based GWAS datasets, however, significant G×E signals have become the rule rather than the exception (11–15). However, these large studies face the opposite problem because they have power to detect subtle technical artifacts (16). For example, heteroskedasticity must be carefully modelled to avoid pervasive false positive G×E in TWAS and heritability estimation (15,17–19).

Here, we explore an underappreciated choice in G×E studies: the scale on which phenotypes are analyzed. For example, because males are typically taller than females, a genetic variant that increases height by 1% will have a greater effect in males on the default centimeter scale. It has long been known that phenotype scale is critical in G×E studies (2,20,21). However, current G×E studies often ignore this decision, implicitly choosing whatever default scale is provided without discussion or robustness analyses.

We establish a framework to distinguish scale-dependent and -independent G×E by identifying the most-additive phenotype scale. We focus on a G×E test based on polygenic scores (PGSxE), which has become the predominant model for G×E in biobanks for its simplicity and power (22). We use a range of phenotypes in UK Biobank to understand which G×E analyses and which phenotypes are liable to scale-dependent bias. We then characterize the impact of phenotype scale on GWAS and PRS, which is substantial for some phenotypes. We conclude by discussing best practices for phenotype scaling to prioritize biologically meaningful genetic discoveries when the optimal phenotype scale is unknown.

## Results

### Scale-dependent and -independent statistical interactions in theory

The existence and magnitude of statistical interactions depend on phenotype measurement scale. By “scale,” we refer to monotonic transformations of a nonnegative quantitative phenotype (21,23), which preserve the phenotype’s rank order. For a given dataset, if an interaction is statistically significant on all considered scales, we call it scale-independent; otherwise, we call it scale-dependent. Our definition explicitly depends on power because we are focused on the new interactions emerging from biobank-scale data.

Any non-linear scale transformation on an additive phenotype will induce interactions with sufficient power (**Note S2**). But, perhaps surprisingly, scale-independent interactions exist. A simple example is the exclusive or (XOR), where a genetic factor, G, increases the phenotype in one environment but decreases it in another; regardless of the phenotype scale, the sign of G’s expected effect will flip based on E (**Note S1.1**). Generalizing this idea, we prove that G×E is scale-independent for Gaussian G and E if the sign of E determines the sign of G’s effect (**Note S1.2**); however, even in this simple case, scale-independence is not robust to outliers (**Note S1.3**). The former conclusion is consistent with prior work relating the sign of G’s effect to scale-independent G×E (24,25) or “crossover” in reaction norm models (26).

### Additive genetic scales can be recovered in simulations

We focus on a simple approach to study G×E using polygenic scores (PGS), which fits PGS×E interaction to test if the PGS effect depends on E. This models the component of G×E that is correlated with additive genetic effects, with E systematically amplifying or dampening the additive effects (27). We fit this model for a range of scales defined as power transformations, i.e., exponentiating the phenotype by λ (**Fig 1)**. When λ=1, no transformation is applied; an important special case is λ=0, which corresponds to log transformation. We then profile PGSxE p-values across λ from -1 to 2, asking if some λ recovers a truly additive scale.

**Figure 1.**
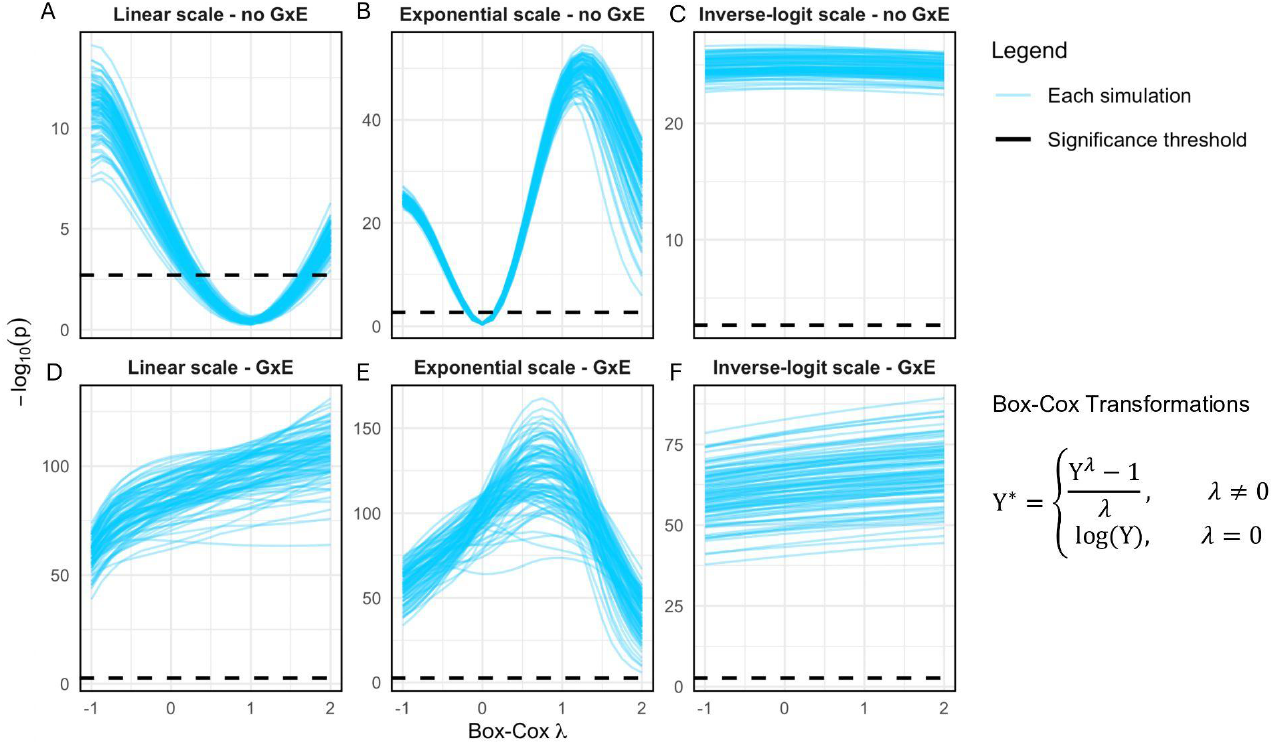
Power transformation removes some forms of scale-dependent G×E in simulations: Each blue line shows the results for testing the interaction between a PGS and an environment, E, averaged over simulation replicates. The Y axis shows the p-value of the PGS×E interaction term and the dotted black line shows p=0.05 after Bonferroni correction. The X axis denotes the different power (or Box-Cox) transformations applied to the phenotype with parameter λ (**Methods)**. Importantly, λ=1 corresponds to no transformation and λ=0 to log transformation.

To test if our approach could distinguish scale-dependent vs -independent PGS×E, we simulated a latent phenotype and then transformed it to mimic an observed phenotype, varying if the latent phenotype is additive (**Fig 1A-C**) or has G×E (**Fig 1D-F**). When the latent scale was directly observed, PGS×E tests were calibrated on the observed scale, as expected; in fact, in this setting, applying a power transformation creates scale-dependent PGS×E (**Fig 1A**). When the phenotype is observed on the exponential scale, however, PGS×E tests were severely inflated; as expected, this was corrected by log transformation (**Fig 1B**). Notably, we do not observe power transformations meaningfully reducing power for when PGS×E is truly present on the latent scale (**Fig 1D-F**). Finally, our approach is unable to remove scale-dependent PGS×E when the inverse-logit scale is observed, as expected because this is not a power transformation (**Fig 1C**).

We also tested the rank-inverse normal transformation (RINT), which scales the phenotype to have approximately Gaussian quantiles. RINT did not depend on the observed scale, as expected, and always gave modest inflation for PGS×E effects (FPR=0.073), even when the additive scale is observed.

In conclusion, separating scale-dependent and -independent G×E is challenging but possible in theory and practice; more importantly, careful consideration of phenotype scale can provide insights into the nature of G×E.

### Scale-dependent G×Sex effects on height are removed by log transformation

We first study height, a classic additive trait in human genetics (28,29). We applied our power transformation procedure to evaluate scale-dependent G×E in ∼300,000 White British individuals in UK Biobank (30) (**Methods**). We tested for PGS×E interactions between 5 environmental variables (E) and 36 polygenic sores that were computed from GWAS that do not include UKB individuals (**Table S3**) (31).

We first study sex as an environment. Surprisingly, despite the limited evidence for sex-specific genetic effects on height (32), we find a highly significant PGS×Sex effect (p=2e-29, **Fig 2A**). Nonetheless, this unexpected effect can be eliminated by log transformation (p=0.09), i.e., power transformation with λ=0. This suggests that genetic effects on height act multiplicatively and that height should be studied on the log scale, which differs from common practice in GWAS (28) but is established in cross-species comparisons (33) and has been shown in humans (34). Across the four other E, we find one additional PGS×E that vanishes on the log scale (alcohol intake frequency), two that were not significant on any scale (smoking status, statin usage), and one that was significant on all tested scales (age, **Fig S1**).

**Figure 2.**
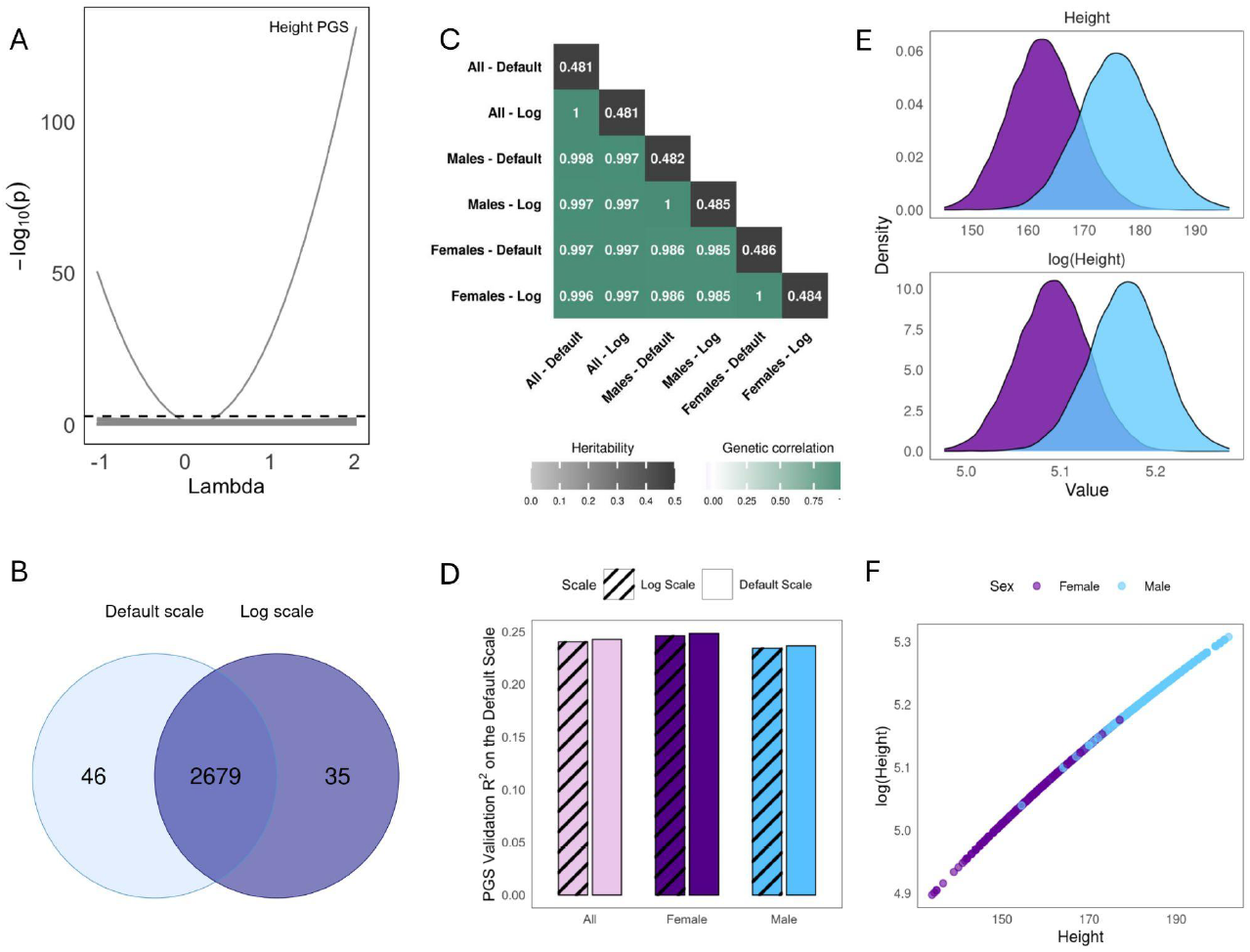
Scale-Dependence in Height: A. Results of the PGS×Sex analysis. Each gray line shows the p-values of the interaction term between a PGS and sex. The dotted black line shows the Bonferroni significance threshold. The height PGS is labeled for clarity. B. Number of GWAS hits when performed on the log scale vs. the default scale. C. Genetic correlations (lower triangle) and heritabilities (diagonal) for GWAS of height, log-height, and sex-specific variants. D. Prediction accuracy for the PGS constructed from the default scale GWAS vs the exponentiated PGS constructed from log-scale GWAS; accuracy is measured by Pearson *R*^*2*^ on the default scale and calculated in either the entire sample or separately in females and males. E. Distribution of height on the default and the log scale, stratified by sex. F. Effect of log transformation on height phenotype values.

We then compared the genetic architecture of height and log-height. We found that their GWAS loci were nearly identical (97.1% overlap, **Fig 2B**; Manhattan and QQ plots in **Fig S2**). Both scales also have similar additive heritabilities and near-perfect genetic correlations with each other and sex-specific GWAS (**Fig 2C**, sex-specific Manhattan plots in **Fig S5**). Furthermore, we found that the PGS constructed from the log and default GWAS gave similar prediction accuracy (**Fig 2D, Methods**). This consistency across scales and sexes is because the log transformation has minimal impact on the distribution of height (**Fig 2E**), providing an almost linear mapping (**Fig 2F**). Overall, the PGS×Sex effect on height illustrates how phenotype scale can dramatically affect G×E tests even when it has negligible impact on additive genetic architecture.

### G×Sex effects on testosterone are largely scale-independent

We next studied an exemplary sex-specific trait to contrast with height: testosterone. Unlike in height, PGS×Sex effects largely remained significant for all tested power transformations (**Fig 3A**), suggesting they are scale-independent. Intuitively, scale-independent interactions are expected because the biology of testosterone qualitatively differs between males and females (35). Supporting this interpretation, all significant PGS×E interactions for the other four environments were scale-dependent (**Fig S3**).

**Figure 3.**
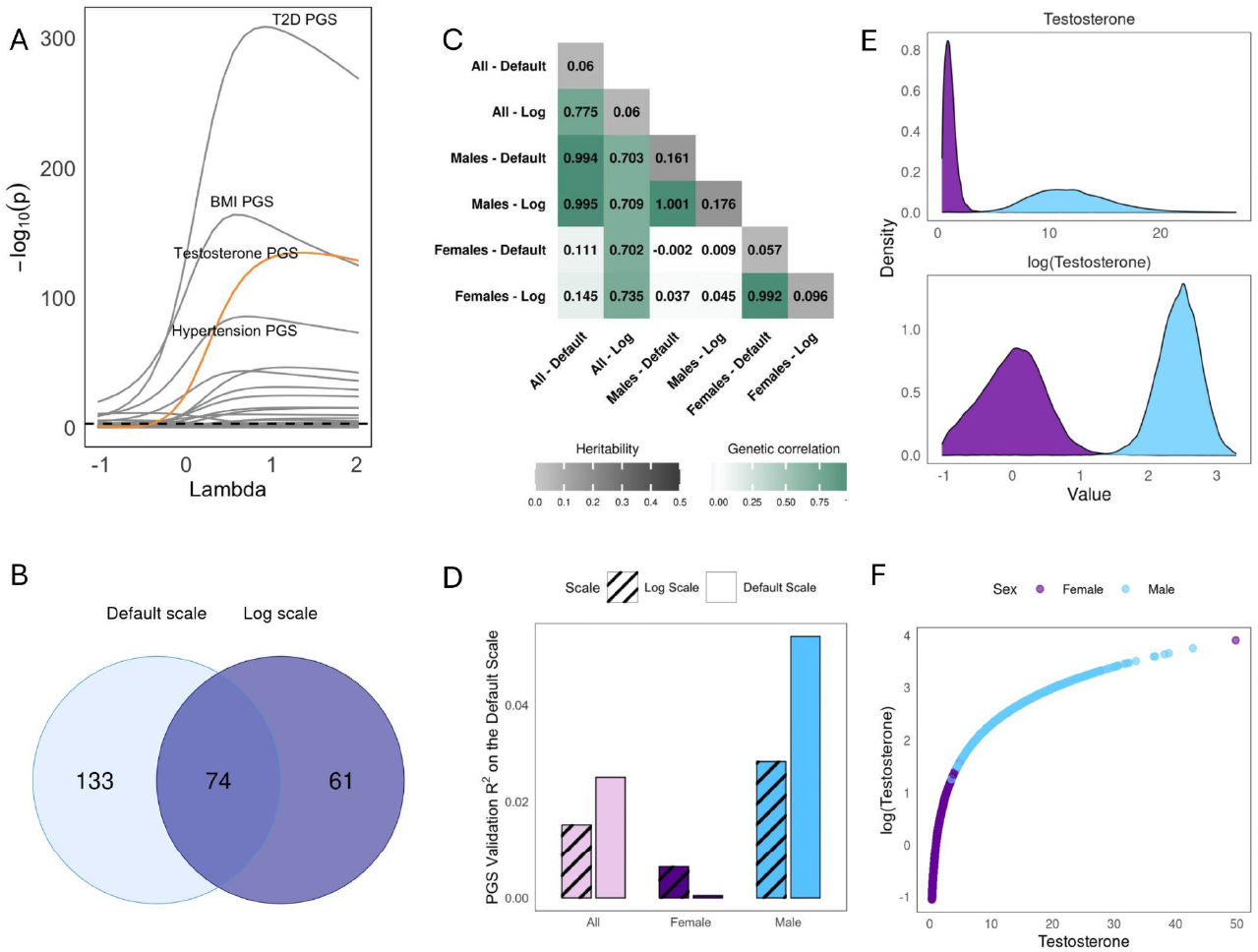
Scale-Dependence in Testosterone: A. Results of the PGS×Sex analysis. Each gray line shows the p-values of the interaction term between a PGS and sex. The dotted black line shows the Bonferroni significance threshold. The four most significant PGS are labeled for clarity. The testosterone PGS (in orange) was constructed in UK Biobank due to lack of external GWAS (**Methods**). B. Number of GWAS hits when performed on the log scale vs. the default scale. C. Genetic correlations (lower triangle) and heritabilities (diagonal) for GWAS of testosterone, log-testosterone, and sex-specific variants. D. Prediction accuracy for the PGS constructed from the default scale GWAS vs the exponentiated PGS constructed from log-scale GWAS; accuracy is measured by Pearson *R*^*2*^ on the default scale and calculated in either the entire sample or separately in females and males. PGS are constructed either from default scale GWAS, or by exponentiating the PGS constructed from log-scale GWAS. E. Distribution of testosterone on the default and the log scale, stratified by sex. F. Effect of log transformation on testosterone phenotype values.

We then compared the testosterone GWAS on the log vs default (nmol/L) scales. Unlike for height, most GWAS hits for testosterone are only detected on one scale (27.6% overlap, **Figs 3B and S4**). We expected this discrepancy arose because males explain the vast majority of phenotypic variance on the default scale but not the log scale (**Fig 3E**). Indeed, we found that the default scale has genetic correlation of 0.99 and 0.15 with the male- and female-specific GWAS (**Fig3C**), respectively, and that its GWAS loci primarily overlap the male-specific GWAS (87.2%, sex-specific Manhattan plots in **Fig S6**). Conversely, the log scale GWAS had much more similar genetic correlations with the sex-specific GWAS (0.70 and 0.74, **Fig3C**). For example, the female-specific effect associated with *FGF9*, which has been linked to gonad development (35–38), is significant on the log scale (p=4.29e-9) but not on the default scale (p=0.13).

As the log scale improves detection of female-specific genetic effects on testosterone, we expected the log-scale PGS would improve prediction. Surprisingly, this reduced Pearson *R*^*2*^ (from 2.50% to 1.52%, **Fig 3D**). However, this is explained by the fact that Pearson *R*^*2*^ is itself scale-dependent: while the Pearson R^*2*^ does indeed dramatically increase for females (13-fold, from 0.05% to 0.65%), it decreases in males (from 5.41% to 2.83%), constituting a net loss because Pearson *R*^*2*^ weights groups by phenotypic variance (**Fig 3E,F**). On the other hand, evaluating PGS *R*^*2*^ on the log scale gives the expected result, where the PGS constructed from the log scale performs better (from 1.02% to 1.74%). This demonstrates how the choice of phenotype scale affects not only PGS but also the relative prioritization of individuals in evaluation metrics. We observe similar results when using Spearman’s rank correlation coefficient to calculate *R*^*2*^ (**Fig S8B**), which is a scale-independent metric but remains sensitive to the GWAS scale used to construct PGS.

Overall, the choice of phenotype scale qualitatively changes inferences on the nature of additive and GxSex effects on testosterone.

### Scale-dependent and -independent G×E are pervasive in biobank-scale data

We expanded our PGS×Sex analysis to 32 quantitative phenotypes (**Table S2**). Across PGS and phenotypes, we find 188 PGS×Sex interactions that are significant on the default scale (15.3% of the total tests, p<0.05/36). Of these, 46.3% become non-significant after power transformation with some λ ∈ [-1,2], and 23.4% are eliminated simply by log transformation (i.e. λ=0, **Fig 4A**). Some phenotypes, such as average fat-free arm mass, mainly have scale-dependent interactions with sex (10/11 vanish on the log scale), while others, such as diastolic blood pressure, only have scale-independent interactions. We find similar patterns for the other four environments, where 7-21% of PGS×E effects are not significant after log transformation (**Fig 4B, S10**). The results are robust when we instead define statistical significance by p<0.05 (**Fig S11**).

**Figure 4.**
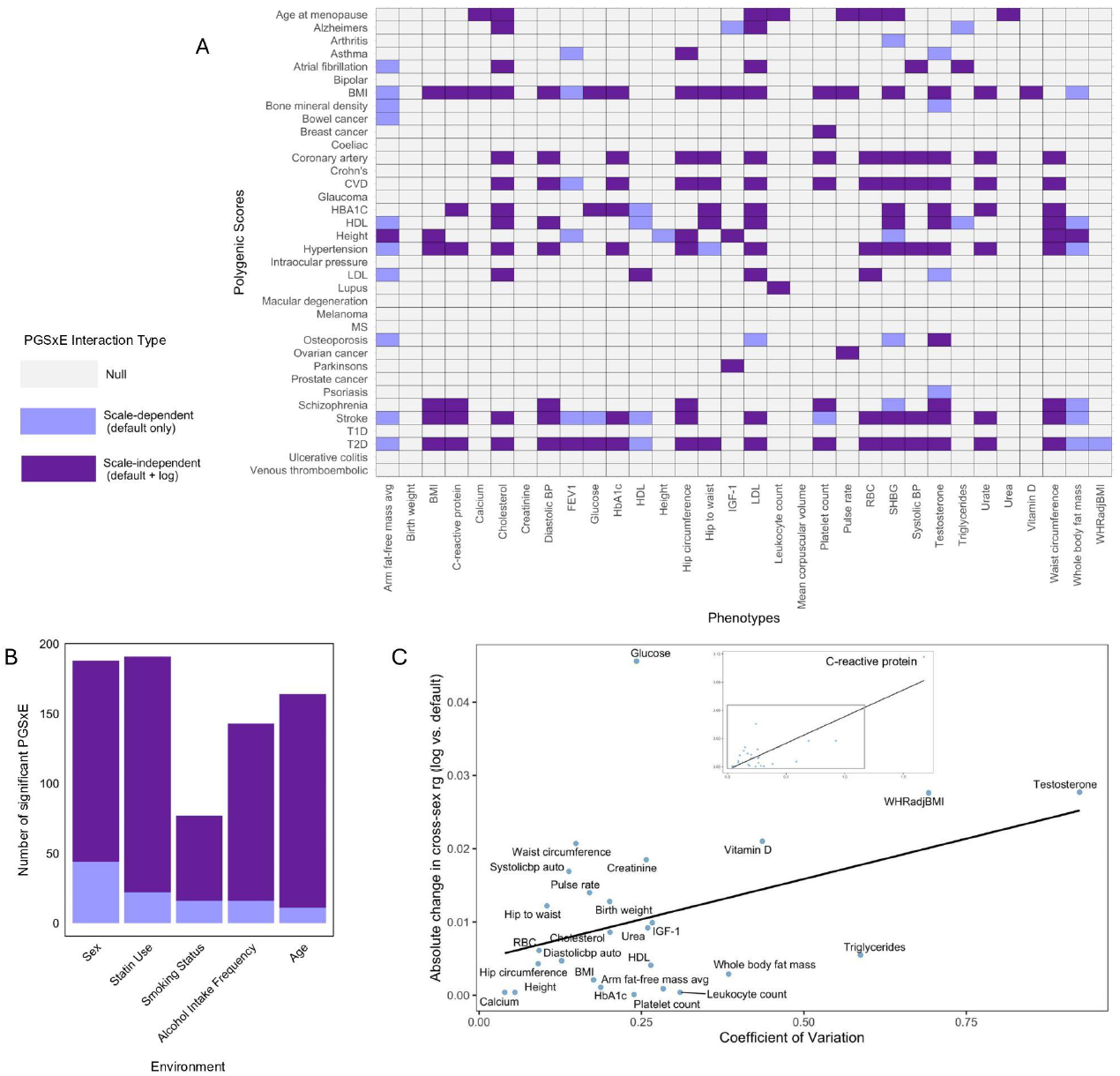
Scale-Dependence in 32 complex traits: A. Results of PGS×Sex analysis across 32 traits. Grey cells show interactions that were insignificant on the default scale; dark purple cells show those that were significant on both the default and the log scale; and light purple cells show those that were significant on the default scale but not on the log scale. B. Number of scale-dependent and scale-independent PGS×E interactions found across all PGS and all phenotypes for each environment. C. Relationship between the coefficient of variation and the absolute value of the change in cross-sex genetic correlation after log scaling the phenotype; relative changes are consistent (**Fig S12.B**)

We also find examples of interactions that cannot be eliminated by power transformation and yet are likely scale-dependent (**Fig S9**). For example, we find complex interactions for biomarkers that are targeted by common drugs, like LDL, which may be eliminated by more sophisticated transformations (17). This highlights how our power transformation approach only provides a lower bound for the severity of scale-dependent bias in G×E.

It is known that scale transformations introduce stronger mean-variance relationships when the coefficient of variation (CV, the ratio of the phenotype’s standard deviation to its mean) is higher (23,39), as we observed in our data (**Fig S12.A**). To ask how CV impacts interactions, we compared cross-sex genetic correlations on the default vs log scale. Across phenotypes, we find a significant relationship between CV and the change in cross-sex genetic correlation after log scaling (p=1.2e7, **Fig 4C, Fig S12.B**). We also find a significant relationship between CV and the change in *R*^*2*^ from constructing the PGS on the default vs log scale (p=1.08e-0.2, **Fig S12.C**). We conclude that the coefficient of variation is one ingredient determining the robustness of a phenotype’s genetic architecture to scale transformation.

## Discussion

We found that choice of phenotype scale can qualitatively change inferences about the existence and nature of gene-environment interactions (G×E) in complex human traits. Although scale-dependence is a textbook concern, its dramatic impact on modern G×E studies is underappreciated. This especially applies to PGS×E models, which are increasingly common (22). Concretely, across 32 phenotypes in UK Biobank, we find that 23.4% of significant PGS×Sex vanish after simply log transforming. In some cases, we also find that GWAS loci and PGS construction and evaluation depend heavily on scale. In general, we find that phenotype scale is more important for phenotypes with larger coefficients of variation (CV).

We contrast G×Sex effects at two ends of the phenotype spectrum: height and testosterone. In height, which is generally considered additive (40), we found highly significant PGS×Sex effects that vanish after log transformation. Nonetheless, its GWAS and PGS are negligibly impacted by choice of scale, partly because height has a low CV. By contrast, in testosterone, which has sex-specific biology (35), we found that G×Sex interactions could not be eliminated by power transformation. Yet still, we found that scale impacts biological discoveries: the log scale GWAS found female-specific effects that are missed by the default scale, such as the *FGF9* effect on gonad development.

Phenotype scale can also impact PGS construction and evaluation. When evaluating Pearson *R*^*2*^ on the default scale, the optimal scale for PGS construction (log or default) varied across phenotypes (**Fig S8.A**), which also held for the scale-independent Spearman *R*^*2*^ (**Fig S8.B**). By contrast, the log-constructed PGS was generally superior when evaluating Pearson *R*^*2*^ on the log scale (**Fig S8.C**). Interestingly, we found that PGS evaluation metrics weigh individuals differently depending on scale. For example, evaluating Pearson *R*^*2*^ for testosterone on the default scale is essentially equivalent to ignoring females. Thus, seemingly subtle modifications to phenotype scale can dramatically impact PGS accuracy and equity.

Across phenotypes, we found that significant PGS×Sex interactions often vanish after power transformation. Many studies have interpreted PGS×E signals as the environment mechanistically amplifying or buffering additive genetic effects (27,41–49). However, amplification signals are expected for log-additive phenotypes; in fact, they are equivalent to first order for binary E (**Note S2.1**). Thus, measurement scale is a major challenge in identifying meaningful forms of amplification.

Other interaction tests are also affected by phenotype scale. For instance, epistasis (G×G) studies in deep mutational scans historically used statistical tests prone to scale artifacts (41,50–52). Phenotype scale is less relevant to epistasis in complex traits because SNP effects are typically weak (53) and PGS are themselves additive (49); however, scale is likely relevant to epistasis between large-effect variants or burden scores. Phenotype scale also applies to binary phenotypes in the form of link function choice, such as probit vs logit (**Fig S13**, (54). More broadly, phenotype scale is liable to confound any non-additive statistical test, such as quantile-dependent effects (55,56), effects on variance (57,58), or machine learning models (57,58).

What scale should be used in practice? It depends on the goal. Ideally, the scale is carefully chosen based on domain expertise, or prespecified goals like maximizing PRS *R*^*2*^ on a clinically-relevant scale (59). Nonetheless, even in this case, prediction may be improved by performing GWAS on a different scale and then transforming back (**Fig S8.A**).

Statistically, the scale can be chosen for parsimony. Early work in quantitative genetics suggests that optimal scales could be defined as those that minimize heteroskedasticity, epistasis, dominance effects, or non-Gaussianity (2,20,21,23). In the context of G×E studies, we propose choosing phenotype scale to minimize G×E signals, corresponding to the null hypothesis that the phenotype is additive on *some* scale. This is natural because significant G×E is expected when an additive phenotype is tested on a non-additive scale. We generally found that RINT eliminated extreme false positives, though it was always slightly inflated and cannot be easily interpreted.

It is much more challenging to define the optimal scale for the goal of learning biology. The link between statistical and biological interaction is tenuous in both directions: biologically meaningful G×E can be statistically insignificant, and biologically-meaningless G×E is expected under many forms of model misspecification, such as an incorrect phenotype scale. Further, scale-dependent G×E can be informative: for example, the observation that G×E effects on height disappear on the log scale tells us that effects on height are generally multiplicative (34), and accounting for scale-dependent interactions can reveal more meaningful interactions that merit further study (41–47). Overall, scale-independence of GxE is neither necessary nor sufficient for biological insight.

Our study is an early effort to characterize the impact of phenotype scaling on biobank-based genetic studies. We focused on power transformations for simplicity, but richer functional forms should be considered in the future (60). It is likely that scale transformation has the greatest impact on the tails of the distribution, which are not the focus of our study but may be important clinically (56,59,61). Further work should also examine the impact of scale on constructed phenotypes, such as BMI or waist-to-hip ratio. In summary, phenotype scale is a critical parameter in biobank-based genetic studies, especially PGSxE, that should be carefully selected, reported, and discussed.

## Methods

### Simulations

We simulated traits using N=100,000, m=25 and *h*^*2*^=0.3 under the following model:

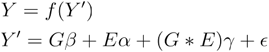

Here, *G* is an *n* x *m* matrix of polygenic scores, *E* is a vector of length of *n* environment values and *ϵ* ∼ *N* (0, 1) is a vector of individual-level noise.*Y*^′^ is the phenotype on the true scale, while *Y* is the phenotype on the observed scale, which was generated using one of three different scaling functions: *f* (*x*) = *x* , *f* (*x*) = *exp* (*x*) , and *f* (*x*).= inverse logit (*x*). We performed 100 simulations using γ =0 and 100 simulations using γ =0.1. The variance explained by the environment was 0.2.

We then try to recover the additive scale, while only knowing *Y,E* and *G*. To this end, we fit a linear model with an interaction term to 25 scaled versions of *Y* using Box-Cox transformations for λ between -1 and 2. These are defined as

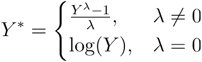

for *Y*^′^ > 0. Additionally, we evaluated rank-inverse normal transformation (RINT) which transforms the quantile of the phenotype to match the quantiles of a Gaussian distribution (62). We determined the significance threshold using Bonferroni correction.

### UK Biobank Data

The UK Biobank is a publicly available prospective dataset from the United Kingdom that contains genetic and deep phenotypic data for participants aged 40-69 (30). In this study, we selected 32 phenotypes to be analyzed, which are listed in supplementary table 2. We manually constructed hip to waist ratio adjusted for BMI, as defined by (42), hip to waist ratio, and average arm fat-free mass (the mean between the right and left arms).

We excluded individuals who had missing data for the phenotype, environment, or covariates in any given analysis. We also excluded outlier individuals whose phenotype value fell in the top or bottom 0.01% of the distribution. Analyses were performed on “White British” individuals, except PGS *R*^*2*^ evaluation was performed on “White European” individuals (30,63). We also excluded one individual from each pair of 3rd degree or closer relatives and individuals with missing genotype call rates greater than 0.01.

### Environment Data

We used five different environmental contexts in our analyses, which are described in **Table S1**. Statin usage was defined using labels defined by (17), which classified individuals as either users or non-users of the drug. Smoking status was modelled as a three-level categorical factor: never smoker, previous smoker and current smoker. Alcohol intake frequency was modeled as continuous, but the variable contains 6 discrete, ordinal values. The fields used for each of these variables are detailed in supplementary table 1.

### PGS×C models in UKB

We fit the following model, for from -1 to 2:

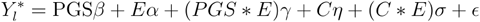

Here 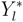 is the Box-Cox (power) transformed value of the phenotype for λ = *l* and *E* is one of the five environmental contexts described above. *PGS* is a standard PGS that was trained on an external dataset, as described by (31) (**Table S3**); the sole exception is the testosterone PGS in **Fig 3.A**, which was not available from external data, so we constructed it in “White British” and tested the interaction in “White European, as outlined below. This PGS is not generally trained on the trait corresponding to 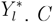 is a matrix of covariates, which contains sex, age, age^*2*^, assessment center and the first 40 principal components. The assessment center is encoded as a categorical factor with 26 different levels. The inclusion of an interaction term between the covariates and the environmental variable (σ) ensures proper adjustment for confounding effects (64). These models were fit in R v4.3.2.

### GWAS and Polygenic Score Construction

We ran GWAS using PLINK2 (v2.00a6LM AVX2 Intel, 4 July 2024) on two versions of each phenotype: once on the baseline scale and once on the log scale (65). For this analysis, we removed SNPs with a missing call rate greater than 0.01, a minor allele frequency smaller than 0.01 or a Hardy-Weinberg equilibrium exact test p-value smaller than 1×10^-6^. This resulted in a total of 6,539,458 SNPs included in the final dataset. We included age, sex and the first ten genetic principal components as covariates. Significant loci were reported with a threshold of 5×10^-8^ after LD clumping with an *r*^*2*^ threshold of 0.1 and a window size of 250kb. We used 90% of the “White British” sample for this analysis.

These GWAS were then used to build PGS, using the same clumping parameters, and these were tested on the remaining 10% of the “White British” cohort. We used a set of p-value thresholds, ranging from 5×10^-8^ to 0.5, and selected the optimal threshold based on prediction performance for the phenotype on the same scale as the GWAS was run, as measured by *R*^*2*^ (generally default-scale Pearson, log-scale Pearson for **Fig S8.C**, and Spearman for **Fig S8.B**).

We then evaluated the PGS in a separate sample, 24,950 “White European” individuals, first on the entire group and then separately on males and females. For scores where the GWAS was trained on the baseline scale, we used the following model.

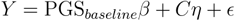

When the GWAS was performed on the log transformed phenotype, we used an alternative model to adjust for this change in scale.

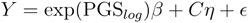

In both models, *C* is a matrix of covariates, containing age, sex and the first 10 genetic principal components. R^*2*^ was used to evaluate each model fit and we compared the two scales using

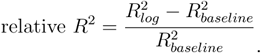

where we emphasize the calculation of correlations is on the default scale. We also performed this procedure using Spearman’s rank correlation to define *R*^*2*^, and the inverse procedure when testing Pearson *R*^*2*^ on the log scale.

### Heritability and Genetic Correlation

We calculated the genetic correlation between sex-specific GWAS and GWAS on the default or log scale. We obtained the GWAS summary statistics on males and females using the procedure outlined above. We then used the package LDSC to obtain heritability estimates for each analysis and estimates of genetic correlation between each pair of analyses (66,67). We filtered SNPs to the HapMap3 set and used LD scores precalculated from the 1000 Genomes European population.

### Identifying Shared and Specific GWAS Hits

To determine the number of shared hits between a pair of GWAS, we first created a third set of p-values, where each SNP was assigned the smallest p-value between those of the two studies of interest. We then clumped this third set, using plink and the parameters outlined above and subset the clumped SNPs to those with a p-value below the threshold of 5e-8. Each of these significant lead hits is then either labeled as shared or only belonging to one of the other two GWAS, depending on whether their p-value was below or above the threshold in each study.

## Code availability

All code used for analyses is available on Github: https://github.com/manuelacostantino1/gxe_scale.

## Acknowledgements

This research has been conducted using the UKB Resource under Application Number 89052. A.D. is supported by R35GM150822. M.C. is supported by Fonds de Recherche du Québec Santé. We thank Carl Veller for helpful feedback on Proposition 1. We thank the participants in UKB for making this study possible. We thank the Center for Research Informatics for providing the computing resources. The Center for Research Informatics is funded by the Biological Sciences Division at the University of Chicago with additional funding provided by the Institute for Translational Medicine, CTSA grant number 2U54TR002389-06 from the National Institutes of Health.

