## Supplementary materials for "Choice of phenotype scale is critical in biobank-based G×E tests"

### 1 Sign-flipping GxE and scale-independent GxE

#### 1.1 The XOR model is a scale-independent form of GxE

We first consider the simplest possible GxE model, where $g$ and $e$ are independent binary variables with mean 0. We assume the phenotype $y$ is fully deterministic and behaves like XOR, with value +1 when the $g$ and $e$ agree and value -1 when they disagree:

$y_i = g_i e_i \iff$

| | $g=-1$ | $g=1$ |
|---|---|---|
| $e=-1$ | 1 | -1 |
| $e=1$ | -1 | 1 |

There is clear GxE on this scale: the main effects of $g$ and $e$ explain 0% phenotypic variation, yet the GxE model explains 100%. But what if we apply a scale transformation $f$? The answer is trivial: only $f(1)$ and $f(-1)$ matter, giving two degrees of freedom in $f$, but neither is identified because mean-centering and variance-standardizing the phenotype comes WLOG. That is, no matter the scale transformation $f$, an identical $y$ is recovered after centering+scaling, so GxE will always explain 100% of the variation in this phenotype even though the additive effects of G and E together explain 0% of the variation.

#### 1.2 A continuous sign-flipping GxE model that is scale-independent

Here, we extend the simplistic XOR idea to a more realistic setting where $g$, $e$, and $y$ are continuous. The key fact we will use about scale transformations is that they preserve signs–a positive effect will be positive on all scales. We will use this to bound GxE based on the estimates in individuals where G has a positive effect vs negative effect. Then, so long as E can distinguish these sets of individuals, the sign of the GxE effect will be identical on all scales, because these sets do not change after scale transformation. This condition on the sign of G's effects is natural because its magnitude can change dramatically by scale transformation, but its sign cannot.

**Proposition 1.** *Assume that $y$ is a deterministic, continuously differentiable function of $g$ and $e$, and that $(g_i, e_i) \overset{\text{iid}}{\sim} \mathcal{N}(0, I_2)$. Assume also that $\frac{dy}{dg} \neq 0$ with nonzero probability. Then $\mathbb{E}\left(\hat{\beta}_{GxE}\right) \neq 0$ for all strictly increasing $f$ if:*

$$sign\left(\frac{dy}{dg}(g,e)\right) = sign(e) \quad \textit{almost everywhere} \qquad (**)$$

*Proof.* First, note that the sign of $g$'s effect is invariant under any increasing $f$:

$$\frac{\partial}{\partial g} f(y) = f'(y)\frac{\partial y}{\partial g} \implies \text{sign}\left(\frac{\partial}{\partial g} f(y)\right) = \text{sign}\left(\frac{\partial y}{\partial g}\right)$$

where sign(0) is taken to be 0.

To simplify bookkeeping of additive effects, we also assume that $f(y)$ is standardized to mean 0 and variance 1. Then the expected GxE regression estimate on scale $f$ is:

$$\begin{aligned}
\beta_{GxE} &= \mathbb{E}_{g,e}\left(gef(y)\right) \\
&= \mathbb{E}_e\left(e\mathbb{E}_g\left(gf(y)|e\right)\right) \\
&= \mathbb{E}_e\left(e\Big[\mathbb{E}_{g|e>0}\left(gf(y)\right) I\{e>0\} + \mathbb{E}_{g|e<0}\left(gf(y)\right) I\{e<0\}\Big]\right) \qquad (\dagger) \\
&= \frac{1}{2}\mathbb{E}_e\left(e\mathbb{E}_{g|e>0}\left(gf(y)\right)|e>0\right) + \frac{1}{2}\mathbb{E}_e\left(e\mathbb{E}_{g|e<0}\left(gf(y)\right)|e<0\right)
\end{aligned}$$

where † breaks the local effect of $g$ given $e$ into the parts where it is positive or negative, and the final step uses our strong assumptions that $g$ and $e$ are independent and that sign $\left(\frac{dy}{dg}\right)$ = sign $(e_i)$.

Then, using Stein's lemma on $g|e$ (which is Gaussian because we assume $(e, g)$ is Gaussian):

$$\mathbb{E}_{g|e>0}\left(gf(y)\right) = \mathbb{V}\left(g|e, e>0\right)\mathbb{E}_{g|e>0}\left(\frac{df(y)}{dg}\right) = \mathbb{E}_{g|e>0}\left(f'(y)\frac{dy}{dg}\right) \geq 0$$

where the final inequality is because $f'$ is always positive and $\frac{dy}{dg} > 0$ whenever $e > 0$. By symmetry, $\mathbb{E}_{g|e<0}\left(gf(y)\right) \leq 0$, and so:

$$\beta_{GxE} = \frac{1}{2}\mathbb{E}_e\left(e\mathbb{E}_{g|e>0}\left(gf(y)\right)|e>0\right) + \frac{1}{2}\mathbb{E}_e\left((-e)\mathbb{E}_{g|e<0}\left(gf(y)\right)|e<0\right) > 0$$

where the inequality in the final line is strict because we assumed $\frac{\partial y}{\partial g}$ is not always 0.

□

On a technical note, our strong assumption (∗∗) is needed to rule out paradoxes where $f$ plucks out a single outlier; if even one data point has the wrong sign for $\frac{\partial y}{\partial g}$, it is possible to magnify its impact to eliminate whatever GxE effect would be estimated from all other data points. This is sketched out in the below counterexample. However, we suspect this condition can be weakened for a stronger GxE test, such as mutual information, as our counterexample relies on an outlier having arbitrarily large influence.

### 1.3 Counterexample: heteroscedasticity frustrates scale-independent GxE

Here, we show how outliers can cause arbitrarily severe biases under extreme scale transformations. In practice, this is important because outliers can create spurious GxE. But in theory, we show here we show how outliers can eliminate GxE that would otherwise be sign-independent.

Suppose the phenotype is generated by the a model that is almost identical to the above section, but now with noise:

$$y = g \circ e + \epsilon; \quad (g_i, e_i) \overset{\text{iid}}{\sim} \mathcal{N}\left(0, I_2\right) \qquad (1)$$

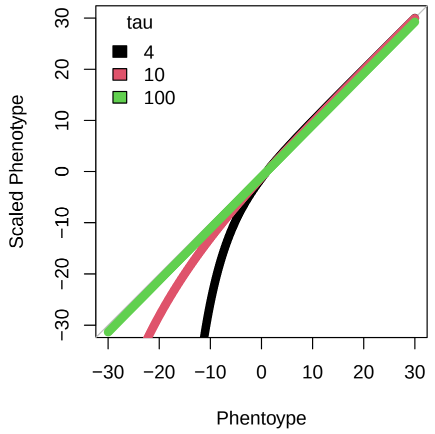


Figure 1: A simple scaling function that exacerbates outliers can arbitrarily impact GxE estimates in a toy model.

Let $\epsilon_i$ by independent mean zero, where all $\epsilon_i$ are distributed between $[-1, 1]$ except that $\epsilon_1 = \{-100, +100\}$ is a mean zero outlier with very high variance.

Without scaling, the expected GxE estimate is 1. But consider a scaling function that essentially acts only by dramatically reducing the negative outlier:

$$f_\tau(y) := xs - \exp\left(-xs + 1\right)/10$$

This transformation will only affect $y_1$, as the other values are close to 0. With 50% chance, this transformation will do nothing (if $\epsilon_1 = 100$); with 50% chance, however, this transformation will replace the expected value $g_1 e_1$ with a much lower value, determined by $\tau$ (shown in Figure 1). In particular, there exists a value of $\tau$ such that $\mathbb{E}\left(\hat{\gamma}_{f_\tau}\right) = 0$, eliminating the GxE.

### 2 Scaling induces PGSxE signals

Assume a standard additive model:

$$y = G + E + \epsilon$$

where $G$ is a polygenic score, $E$ is a univariate environment, $\epsilon$ is iid noise, and all are jointly Gaussian. For simplicity, we again assume $G$ and $E$ are independent and mean zero.

Let $f$ be some monotone increasing function (followed by linear centering+scaling). Applying Stein's lemma twice shows that the GxE estimate has the sign of the average curvature in $f$:

$$\begin{aligned}
\mathbb{E}\left(\hat{\beta}_{G\times E}\right) &= \mathbb{E}\left(f(G + E + \epsilon) * G * E\right) \\
&= \mathbb{E}\left(E\mathbb{E}_G\left(f(G + E + \epsilon) * G\right)\right) \\
&= \mathbb{E}\left(E\left(\sigma_G^2\mathbb{E}_G\left(f'(G + E + \epsilon)\right)\right)\right) \\
&= \sigma_E^2\sigma_G^2\mathbb{E}\left(f''(G + E + \epsilon\right)
\end{aligned}$$

In particular, in the special case where $f$ is convex (or concave) (Sadowski et al. 2025 bioRxiv), then the GxE will always be positive (or negative).

### 2.1 Exponential scaling and a binary environment

In the special case where $f = \exp$, it is clear from above that the interaction between the PGS and environment will be positive. But a simpler direct argument applies in the case where $G$, $E$, and $\epsilon$ are independent and $E$ is binary:

$$\begin{aligned}\mathbb{E}\left(\hat{\beta}_G | E = e\right) &= (e \exp(e))\, \mathbb{E}\left(g_i \exp(g_i)\right) \mathbb{E}\left(\exp(\epsilon)\right) \\ &\equiv e \tau_g \tau_\epsilon\end{aligned}$$

where the last line simplifies by (1) defining $\tau_g := \mathbb{E}\left(g_i \exp(g_i)\right)$ and $\tau_\epsilon := \mathbb{E}\left(\exp(\epsilon)\right)$ and (2) the fact that $E$ and $E \exp(E)$ are equivalent for binary $E$ modulo centering and scaling $E$ (which does not change the GxE sign). Therefore, when additive phenotype with binary $E$ is exponentially transformed, we expect the PGSxE interaction to be significant.

### Supplementary Figure 1: Additional PGS×E Analyses for Height

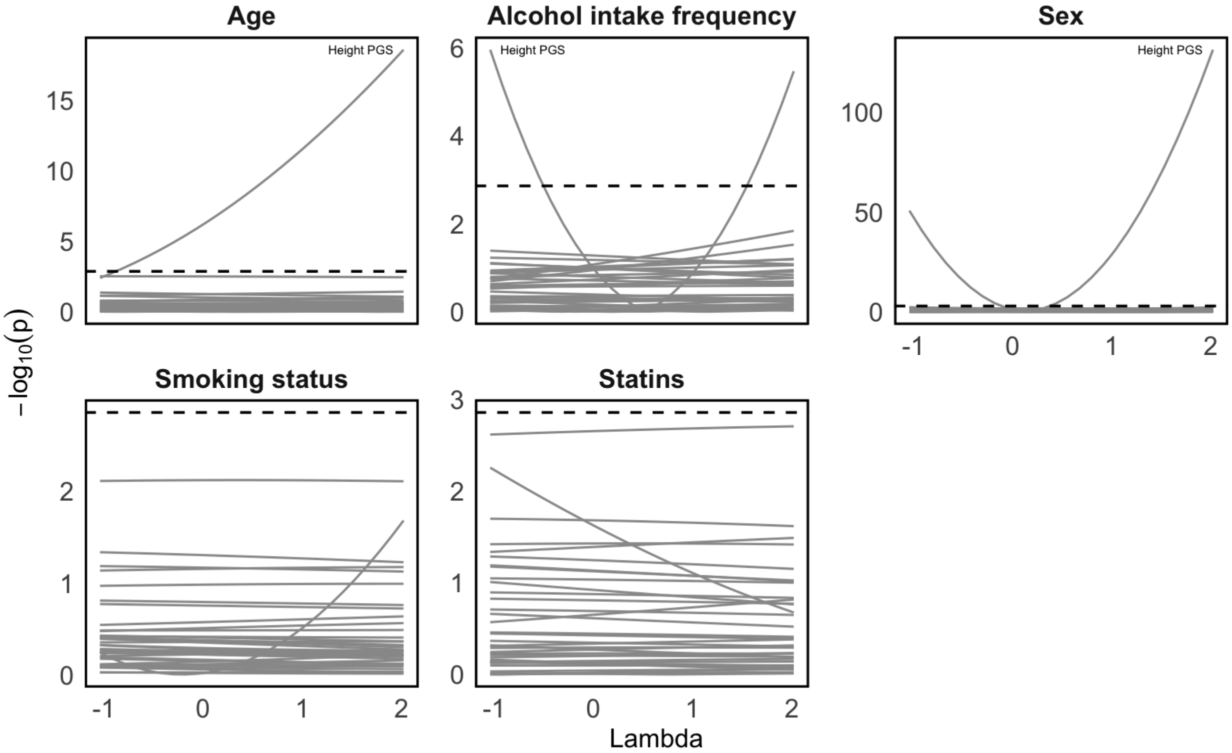

**Supplementary Figure 2: Manhattan and QQ Plots for Height**

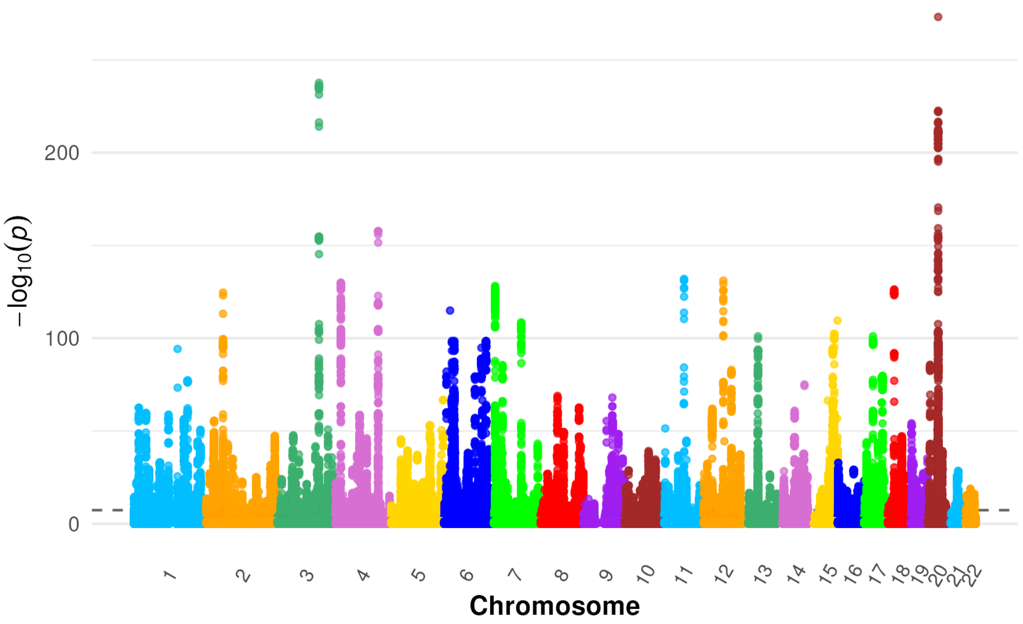


Supplementary figure 2.A: Manhattan plot for height on the default scale. The dotted line is drawn at 5e-8.

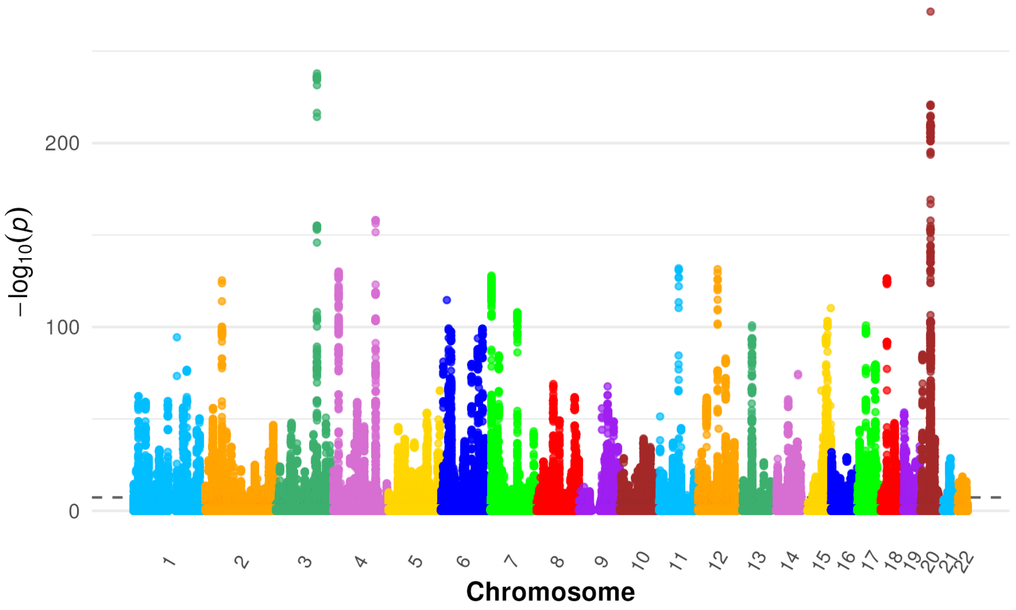


Supplementary figure 2.B: Manhattan plot for height on the log scale. The dotted line is drawn at 5e-8.

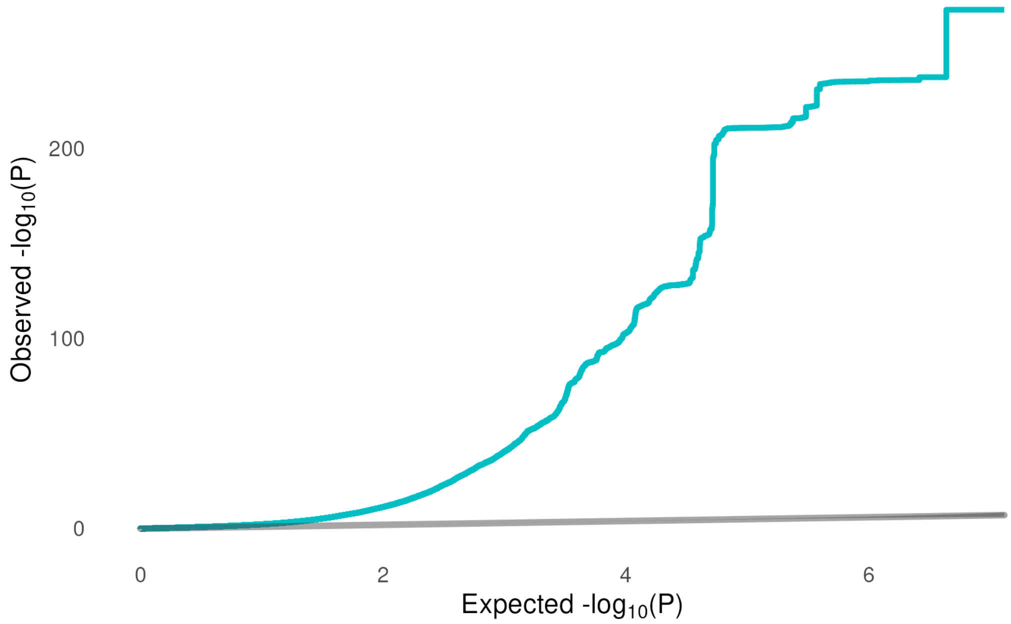


Supplementary figure 2.C: QQ plot for the GWAS of height on the default scale. $\lambda_{GC} = 2.15$

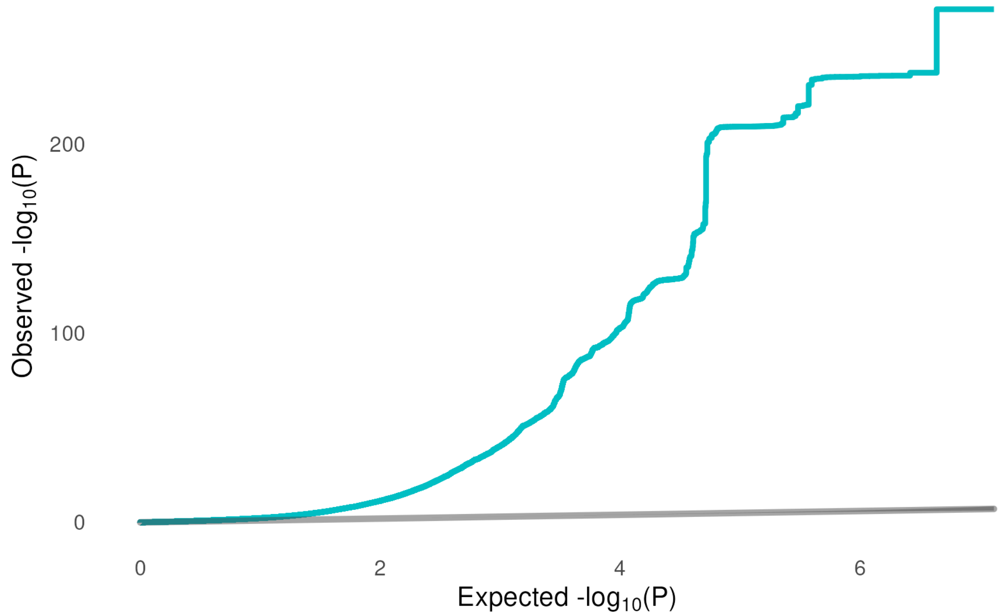


Supplementary figure 2.D: QQ plot for the GWAS of height on the log scale. $\lambda_{GC} = 2.15$

**Supplementary Figure 3: Additional PGS×E Analyses for Testosterone**

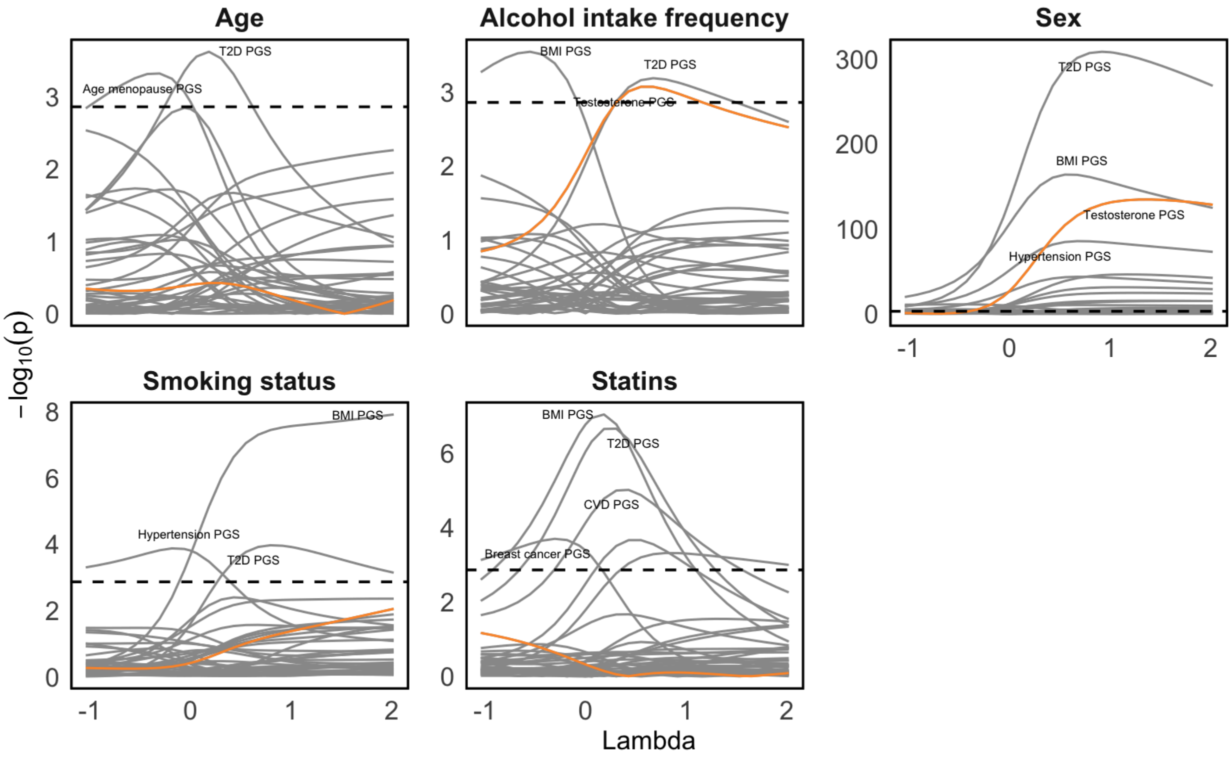

**Supplementary Figure 4: Manhattan and QQ Plots for Testosterone**

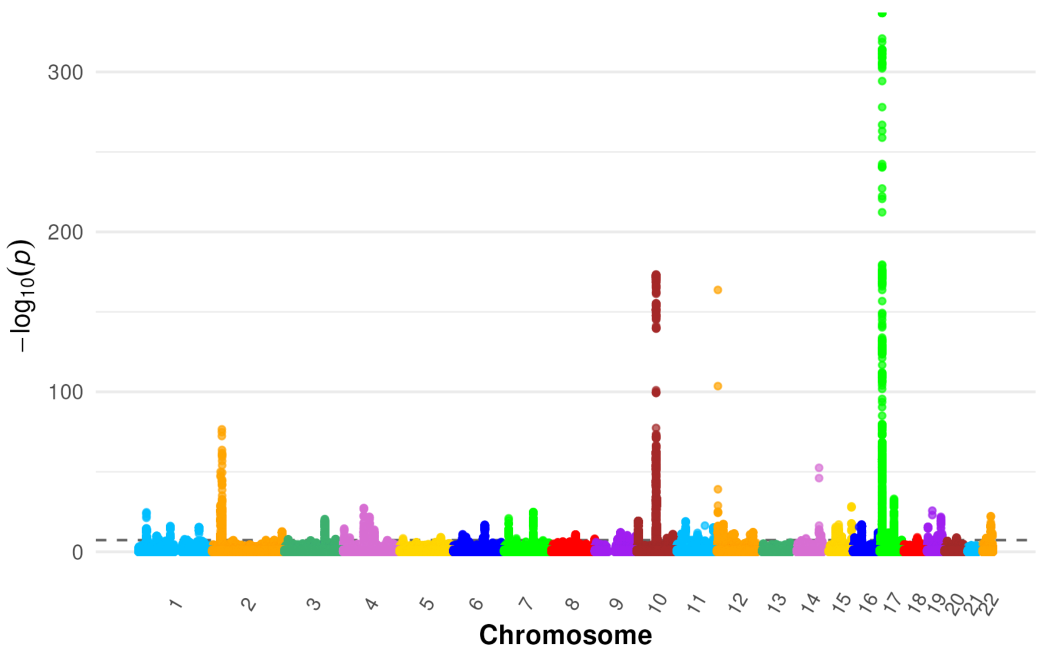


Supplementary figure 4.A: Manhattan plot for testosterone on the default scale. The dotted line is drawn at 5e-8.

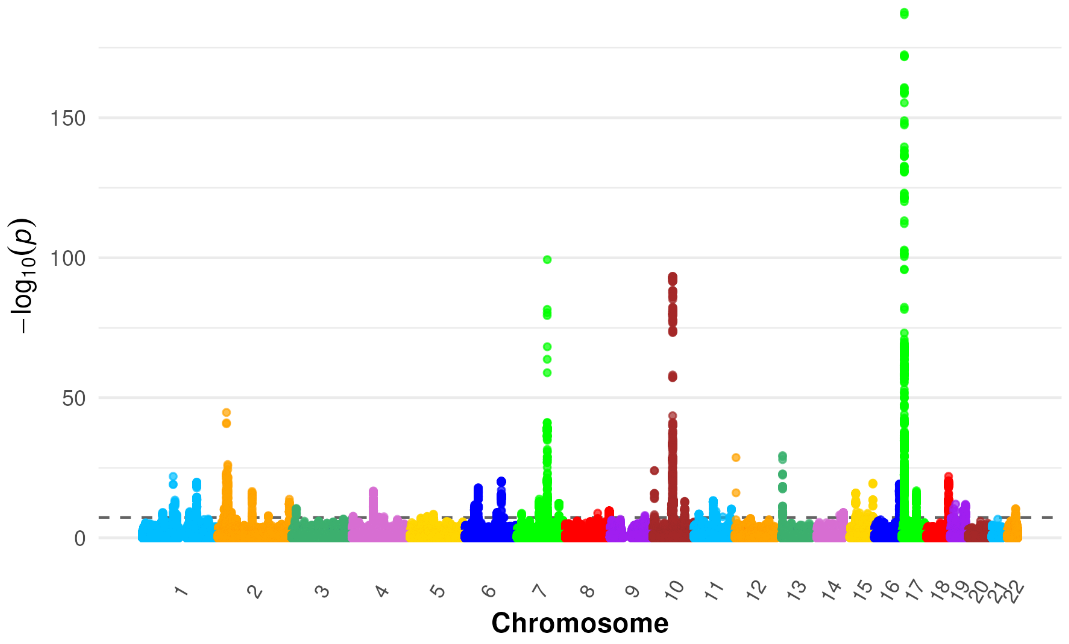


Supplementary figure 4.B: Manhattan plot for testosterone on the log scale. The dotted line is drawn at 5e-8.

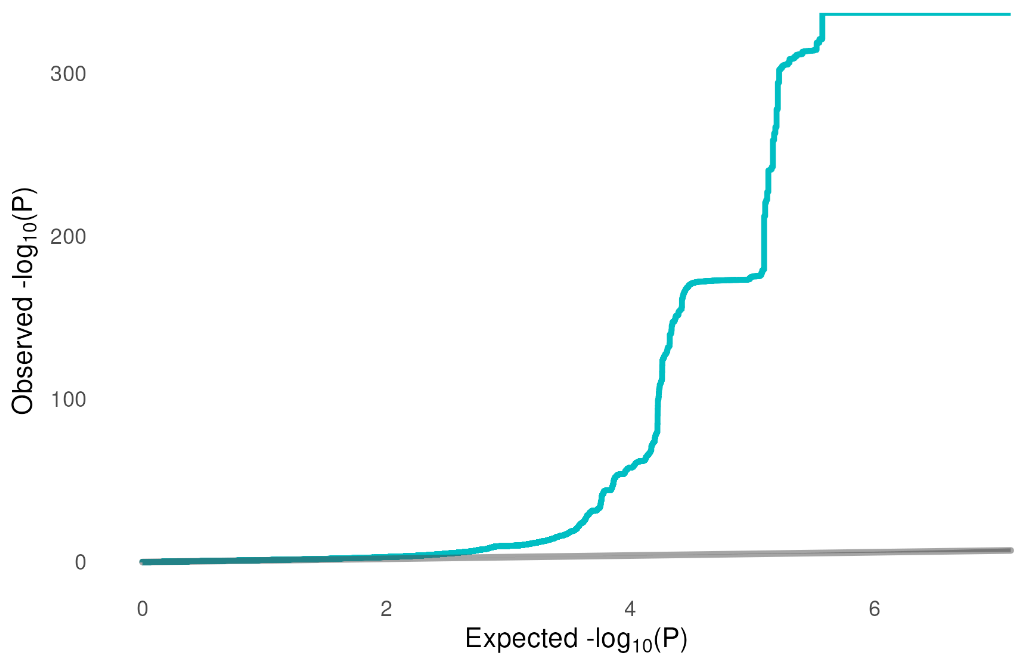


Supplementary figure 4.C: QQ plot for the GWAS of testosterone on the default scale. $\lambda_{GC} = 1.23$

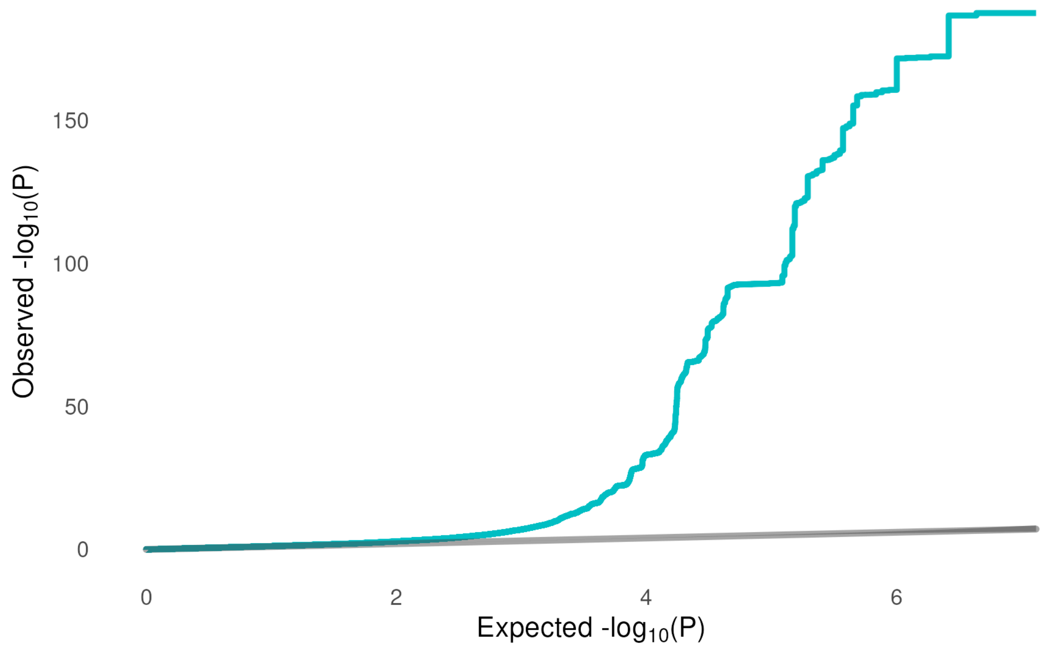


Supplementary figure 4.D: QQ plot for the GWAS of testosterone on the log scale. $\lambda_{GC} = 1.20$

**Supplementary Figure 5: Sex-Specific GWAS for Height**

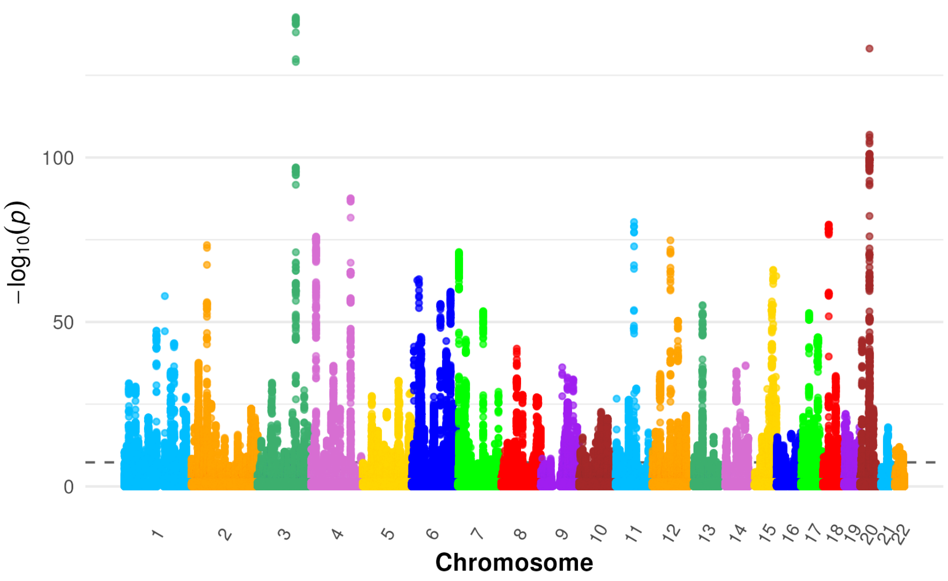


Supplementary figure 5.1: Manhattan plot for GWAS of height in female participants only

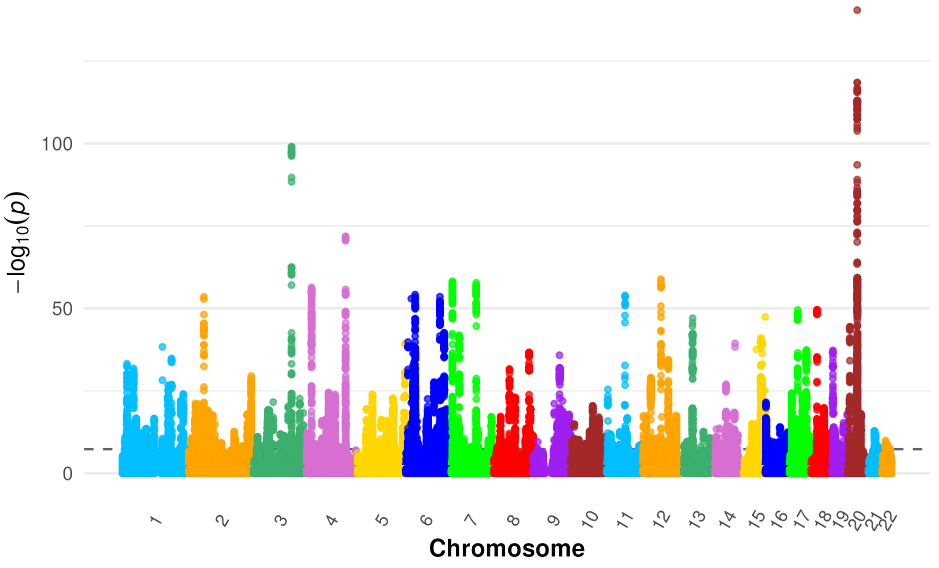


Supplementary figure 5.2: Manhattan plot for GWAS of height in male participants only

### Supplementary Figure 6: Sex-Specific GWAS for Testosterone

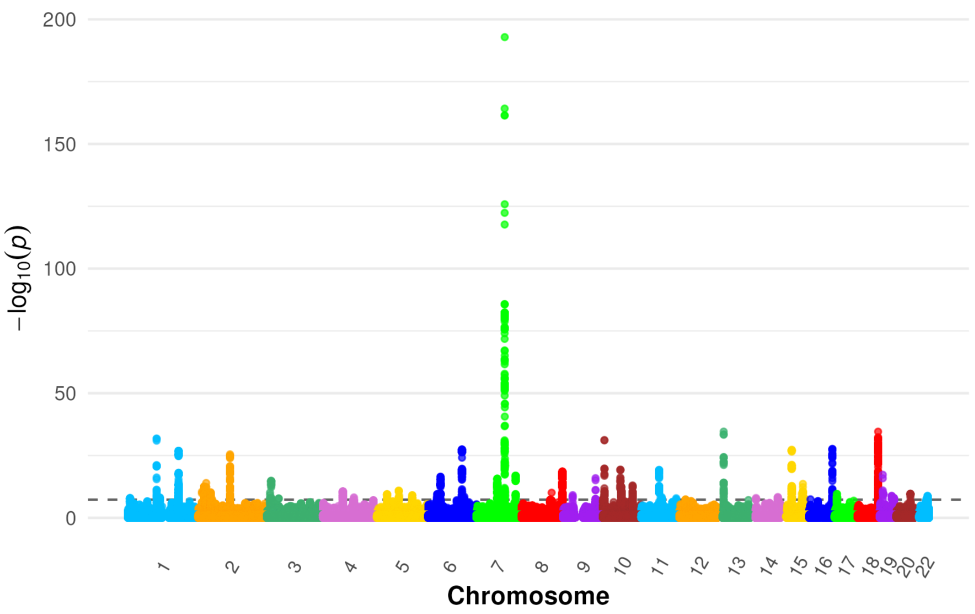


Supplementary figure 6.1: Manhattan plot for GWAS of testosterone in female participants only. Note the significant effect on *FGF9* in chromosome 13 (p=4.29e-9).

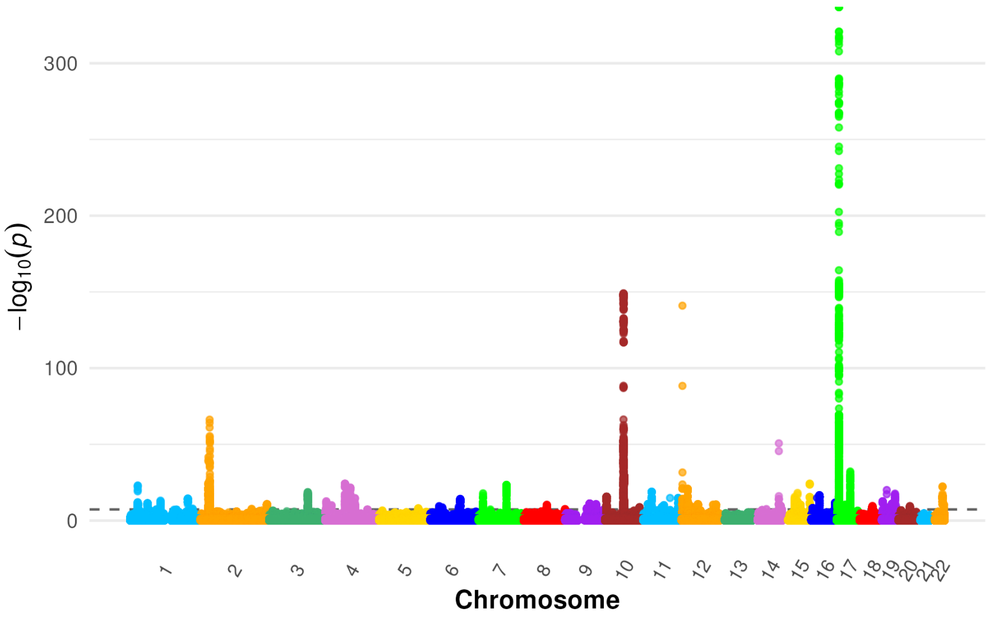


Supplementary figure 6.2: Manhattan plot for GWAS of testosterone in male participants only. Note the insignificant effect on *FGF9* in chromosome 13 (p=0.13)

### Supplementary Figure 7: GWAS of other Phenotypes

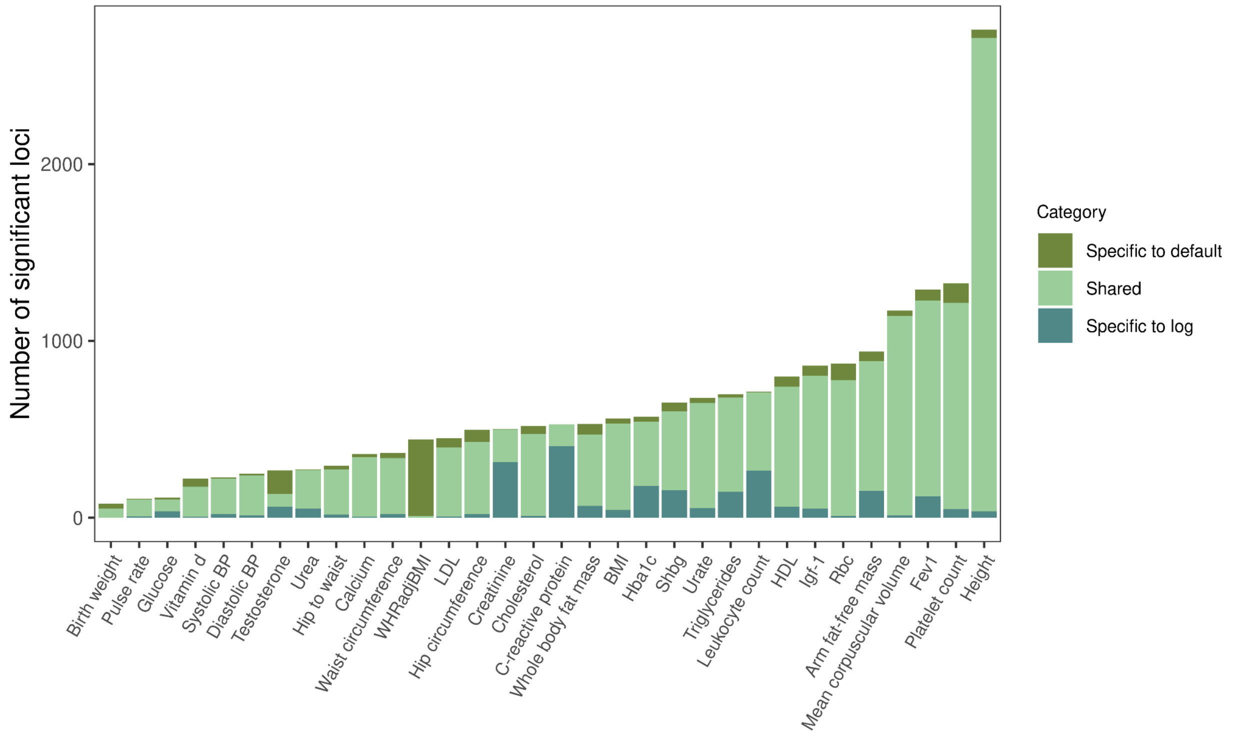

### Supplementary Figure 8: Prediction accuracy of polygenic scores built on the log scale, relative to the default scale

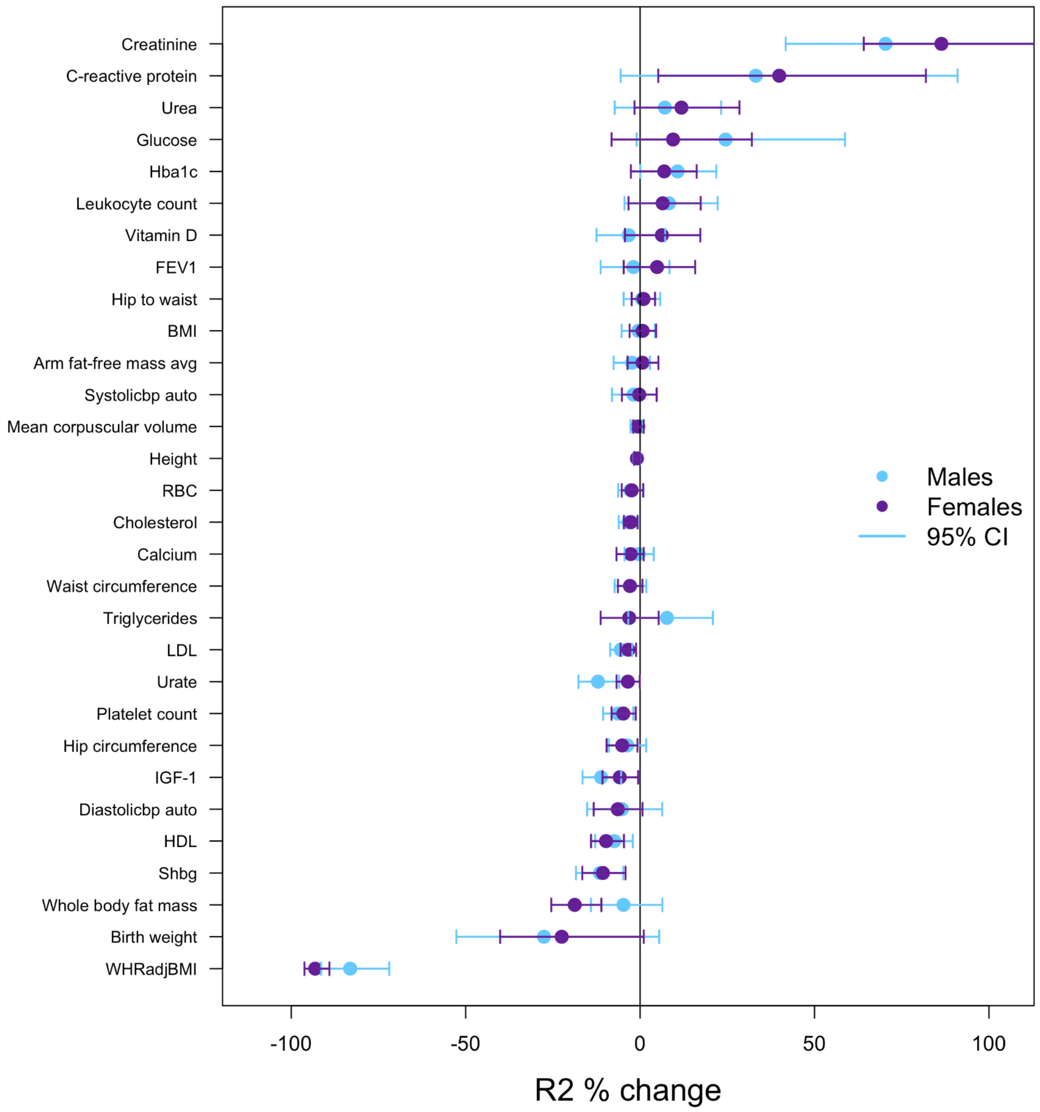


Figure 8.A: Prediction accuracy of polygenic scores built on the log scale, relative to the default scale as measured by Pearson's $R^2$ on the default scale. Negative values indicate increased performance on the default scale and positive values indicate increased performance on the log scale. For testosterone, females see a 1279.56% increase in $R^2$ on the log scale and males see a 47.68% increase on the default scale.

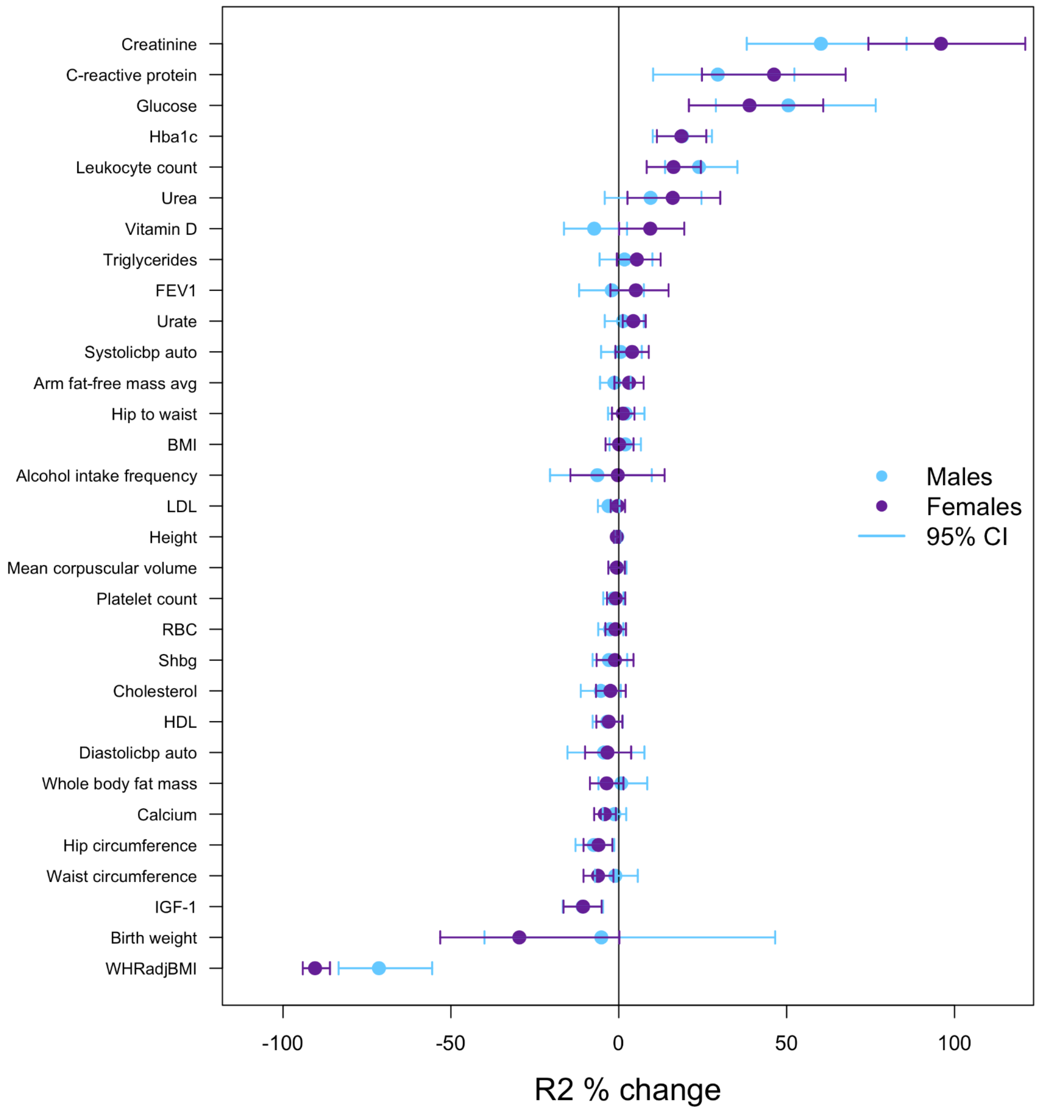


Figure 8.B: Prediction accuracy of polygenic scores built on the log scale, relative to the default scale as measured by Spearman's $R^2$ on the default scale. Negative values indicate increased performance on the default scale and positive values indicate increased performance on the log scale. For testosterone, females see a 2541.03% increase in $R^2$ on the log scale and males see a 46.20% increase on the default scale.

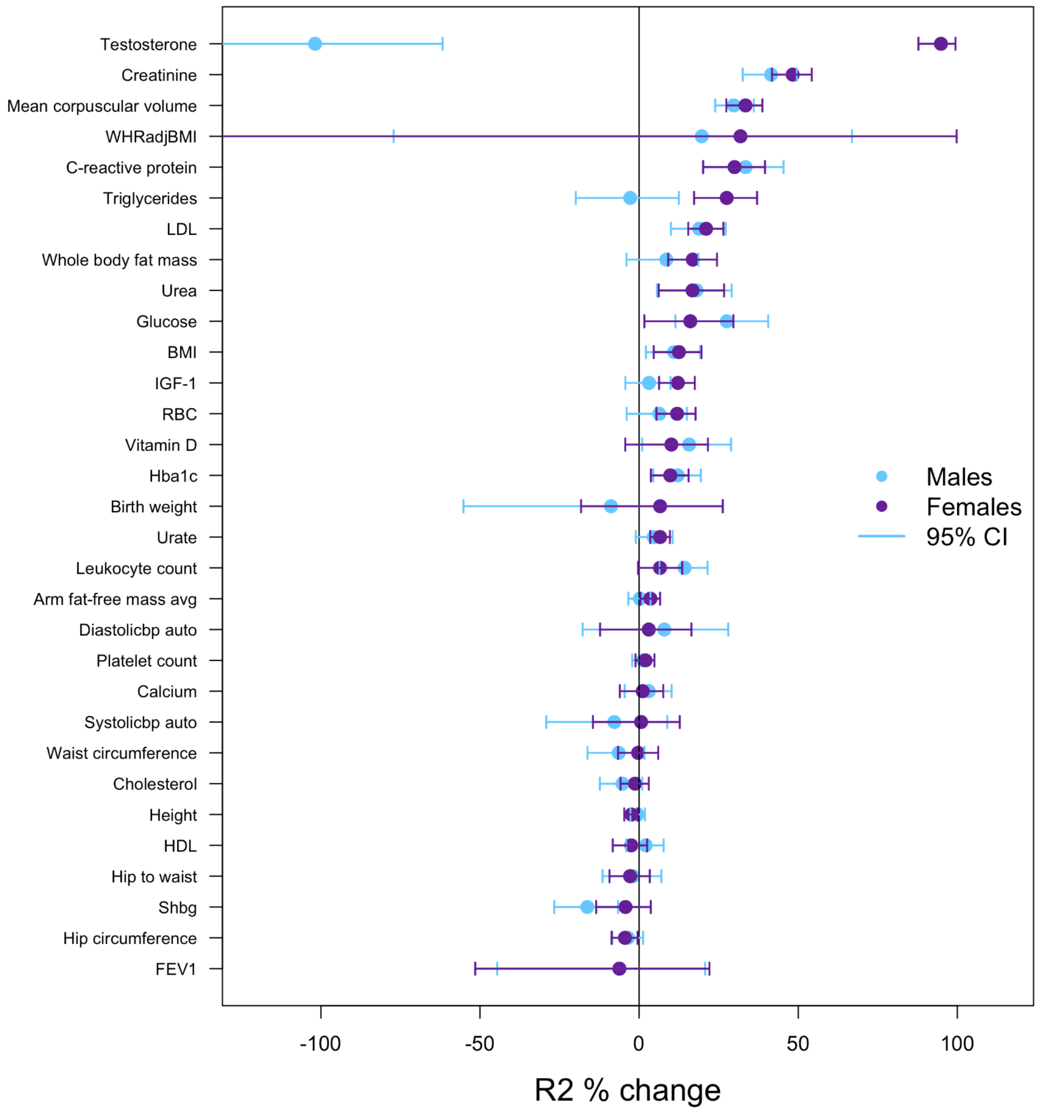


Figure 8.C: Prediction accuracy of polygenic scores built on the log scale, relative to the default scale as measured by Pearson's $R^2$ on the log scale. Negative values indicate increased performance on the default scale and positive values indicate increased performance on the log scale.

**Supplementary Figure 9: LDL and Statin Interaction**

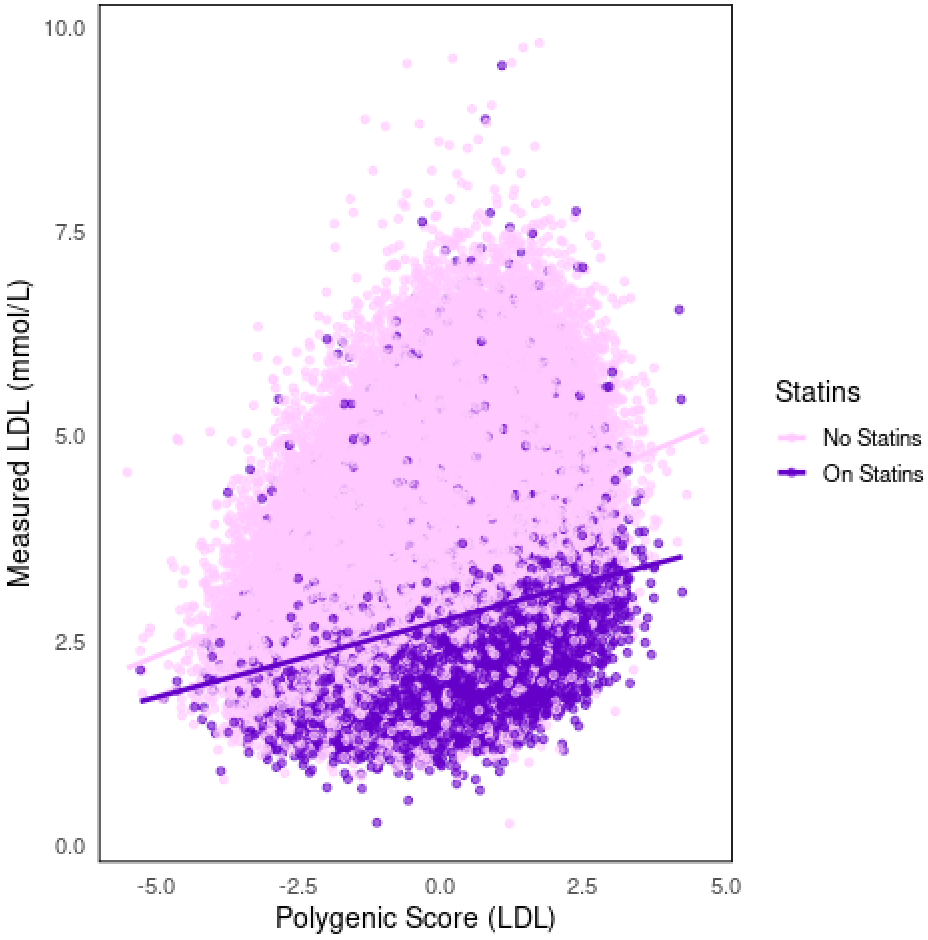

### Supplementary Figure 10: PGS×E Analyses for other Environments

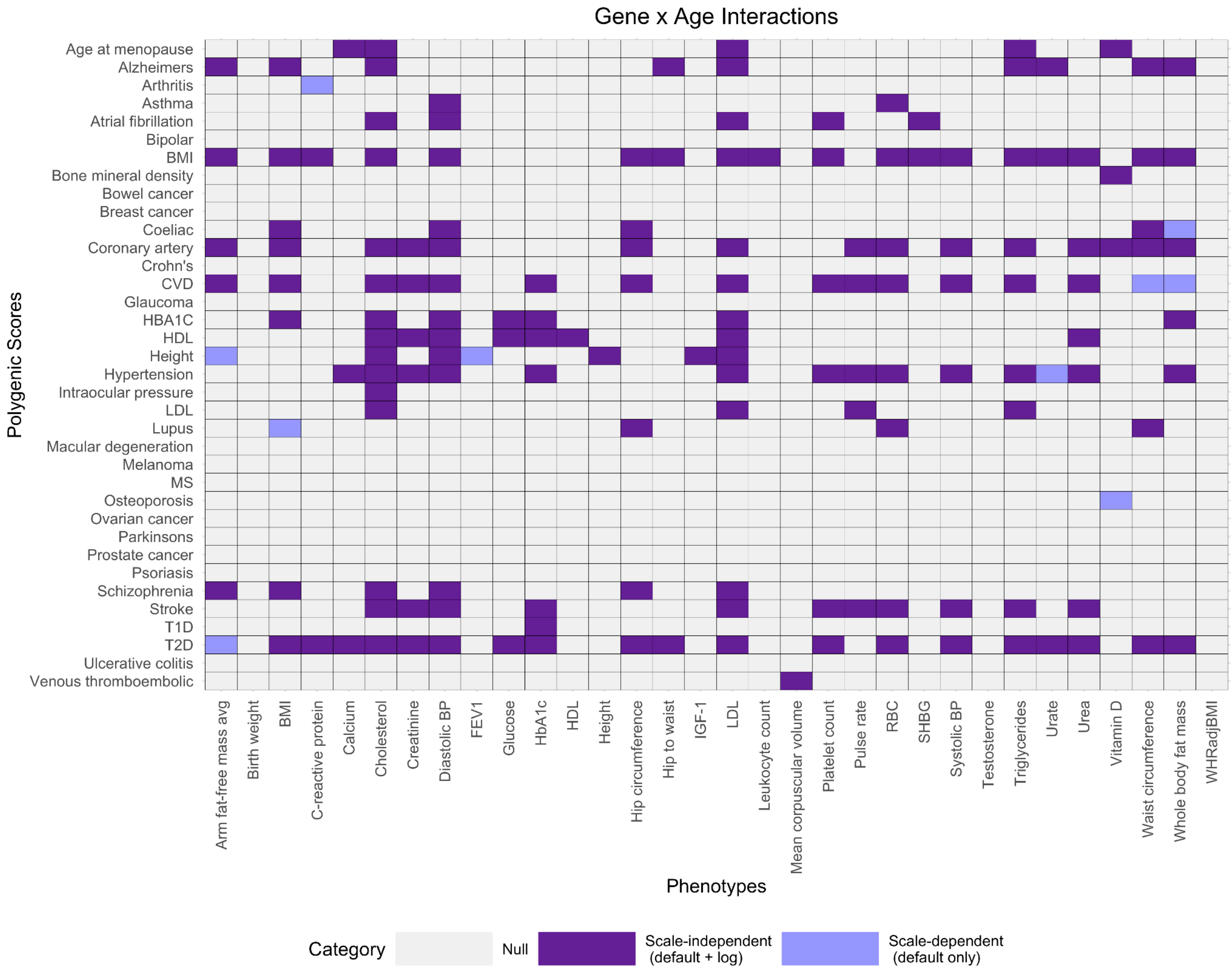

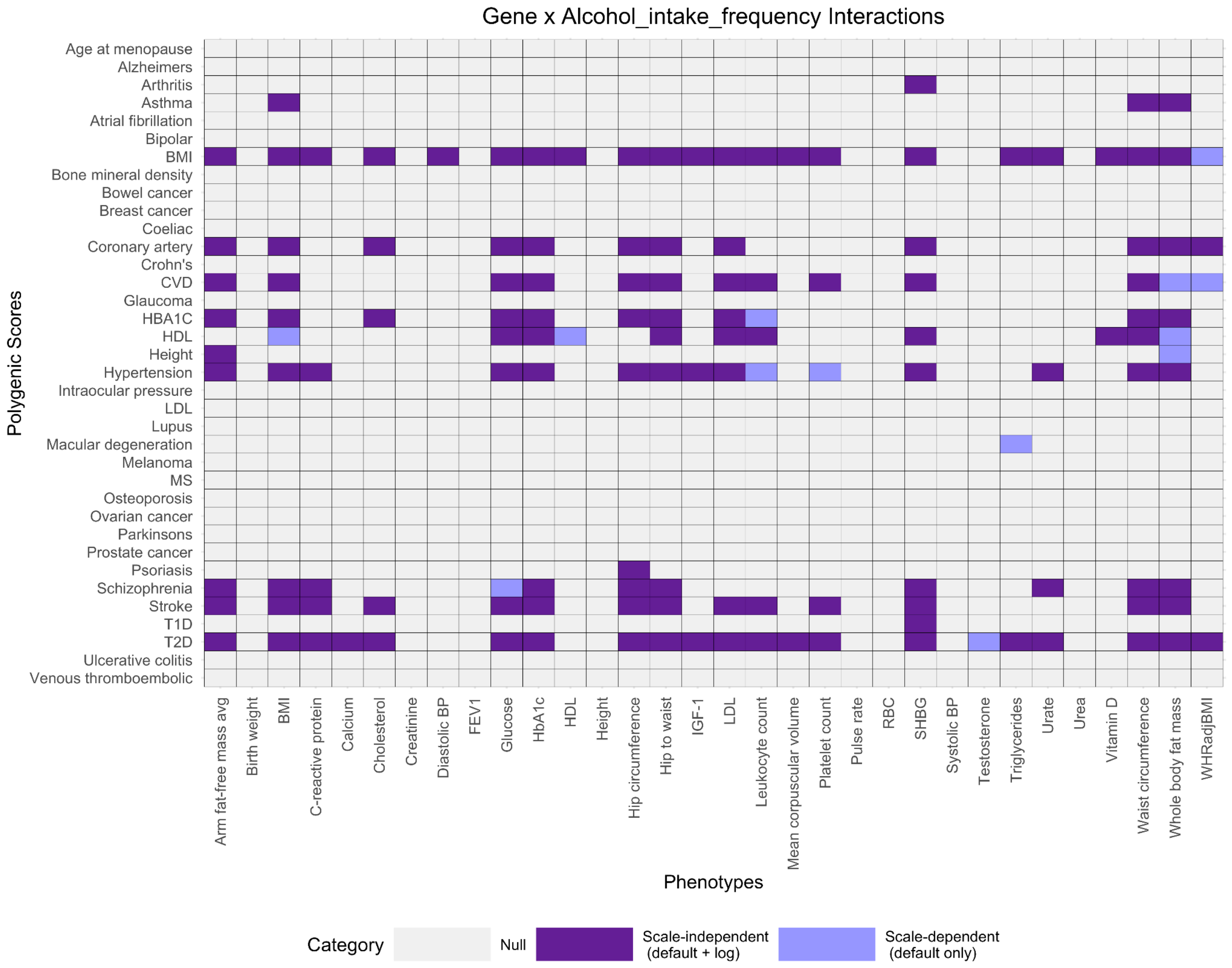
Gene x Alcohol_intake_frequency Interactions
Polygenic Scores
Age at menopause
Alzheimers
Arthritis
Asthma
Atrial fibrillation
Bipolar
BMI
Bone mineral density
Bowel cancer
Breast cancer
Coeliac
Coronary artery
Crohn's
CVD
Glaucoma
HBA1C
HDL
Height
Hypertension
Intraocular pressure
LDL
Lupus
Macular degeneration
Melanoma
MS
Osteoporosis
Ovarian cancer
Parkinsons
Prostate cancer
Psoriasis
Schizophrenia
Stroke
T1D
T2D
Ulcerative colitis
Venous thromboembolic
Arm fat-free mass avg
Birth weight
BMI
C-reactive protein
Calcium
Cholesterol
Creatinine
Diastolic BP
FEV1
Glucose
HbA1c
HDL
Height
Hip circumference
Hip to waist
IGF-1
LDL
Leukocyte count
Mean corpuscular volume
Platelet count
Pulse rate
RBC
SHBG
Systolic BP
Testosterone
Triglycerides
Urate
Urea
Vitamin D
Waist circumference
Whole body fat mass
WHRadjBMI
Phenotypes
Category
Null
Scale-independent (default + log)
Scale-dependent (default only)

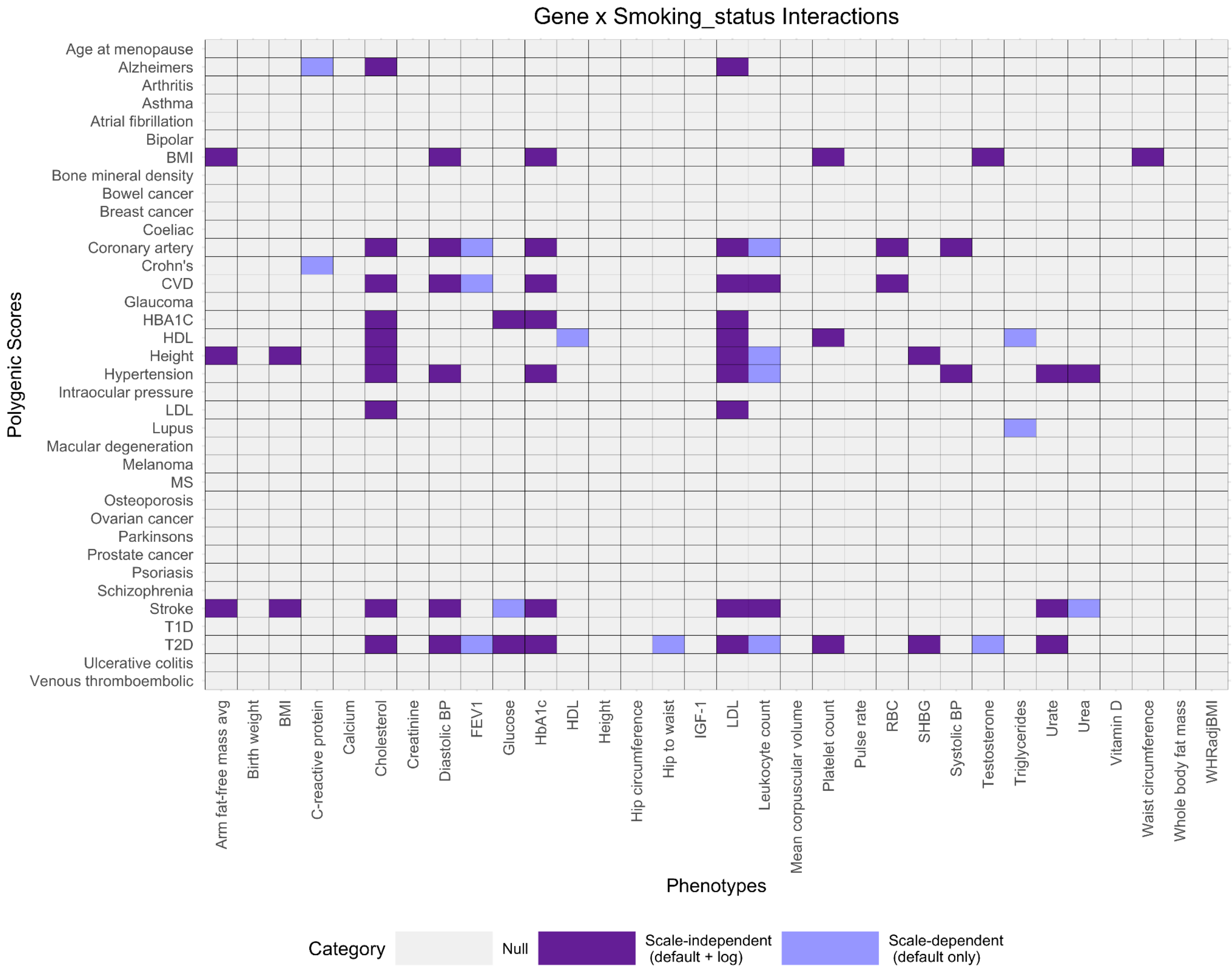

Gene x Smoking_status Interactions
Polygenic Scores
Age at menopause
Alzheimers
Arthritis
Asthma
Atrial fibrillation
Bipolar
BMI
Bone mineral density
Bowel cancer
Breast cancer
Coeliac
Coronary artery
Crohn's
CVD
Glaucoma
HBA1C
HDL
Height
Hypertension
Intraocular pressure
LDL
Lupus
Macular degeneration
Melanoma
MS
Osteoporosis
Ovarian cancer
Parkinsons
Prostate cancer
Psoriasis
Schizophrenia
Stroke
T1D
T2D
Ulcerative colitis
Venous thromboembolic
Arm fat-free mass avg
Birth weight
BMI
C-reactive protein
Calcium
Cholesterol
Creatinine
Diastolic BP
FEV1
Glucose
HbA1c
HDL
Height
Hip circumference
Hip to waist
IGF-1
LDL
Leukocyte count
Mean corpuscular volume
Platelet count
Pulse rate
RBC
SHBG
Systolic BP
Testosterone
Triglycerides
Urate
Urea
Vitamin D
Waist circumference
Whole body fat mass
WHRadjBMI
Phenotypes
Category
Null
Scale-independent (default + log)
Scale-dependent (default only)

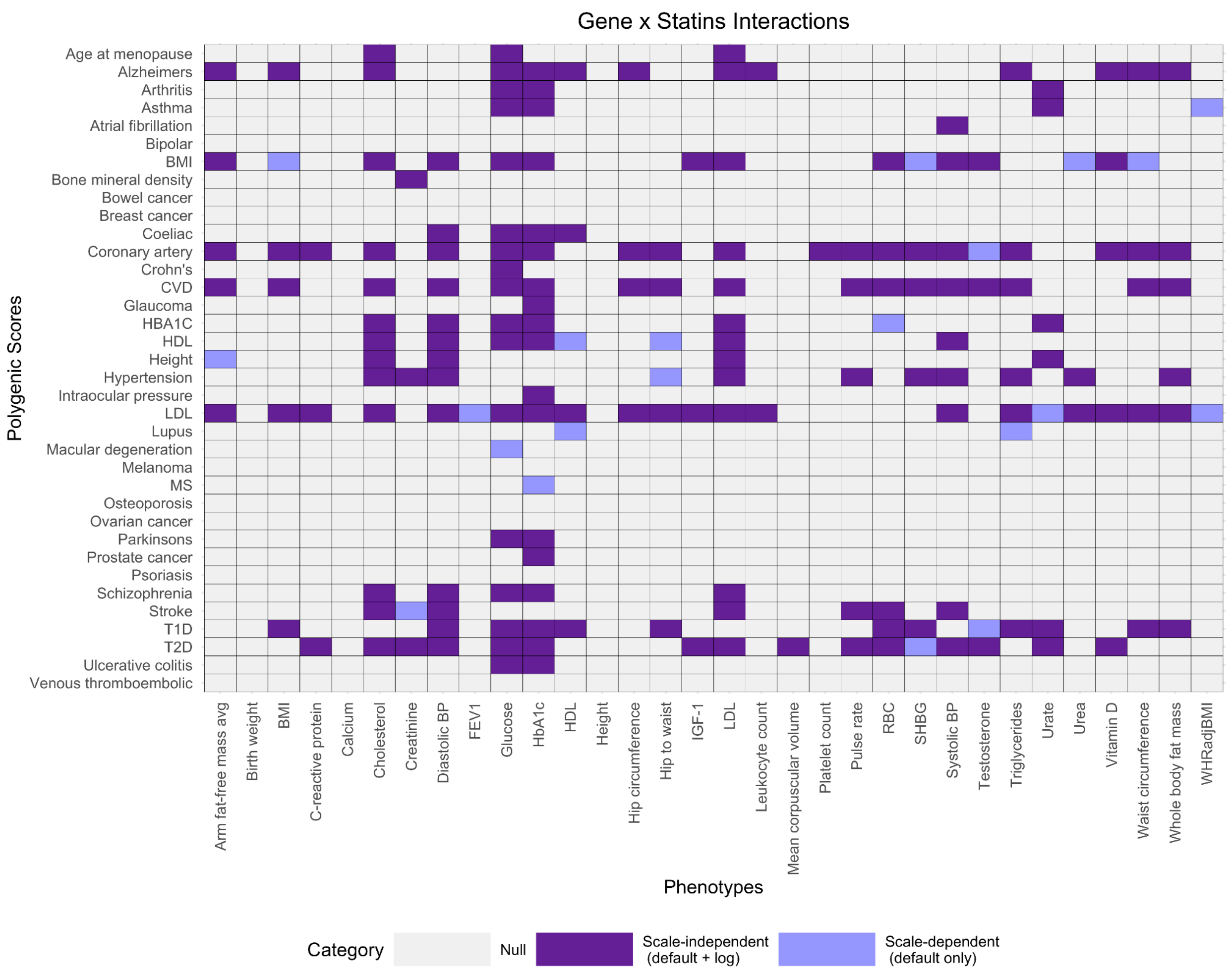

Gene x Statins Interactions
Polygenic Scores
Age at menopause
Alzheimers
Arthritis
Asthma
Atrial fibrillation
Bipolar
BMI
Bone mineral density
Bowel cancer
Breast cancer
Coeliac
Coronary artery
Crohn's
CVD
Glaucoma
HBA1C
HDL
Height
Hypertension
Intraocular pressure
LDL
Lupus
Macular degeneration
Melanoma
MS
Osteoporosis
Ovarian cancer
Parkinsons
Prostate cancer
Psoriasis
Schizophrenia
Stroke
T1D
T2D
Ulcerative colitis
Venous thromboembolic
Arm fat-free mass avg
Birth weight
BMI
C-reactive protein
Calcium
Cholesterol
Creatinine
Diastolic BP
FEV1
Glucose
HbA1c
HDL
Height
Hip circumference
Hip to waist
IGF-1
LDL
Leukocyte count
Mean corpuscular volume
Platelet count
Pulse rate
RBC
SHBG
Systolic BP
Testosterone
Triglycerides
Urate
Urea
Vitamin D
Waist circumference
Whole body fat mass
WHRadjBMI
Phenotypes
Category
Null
Scale-independent (default + log)
Scale-dependent (default only)

**Supplementary Figure 11: Scale-dependent and -Independent Interactions using p=0.05**

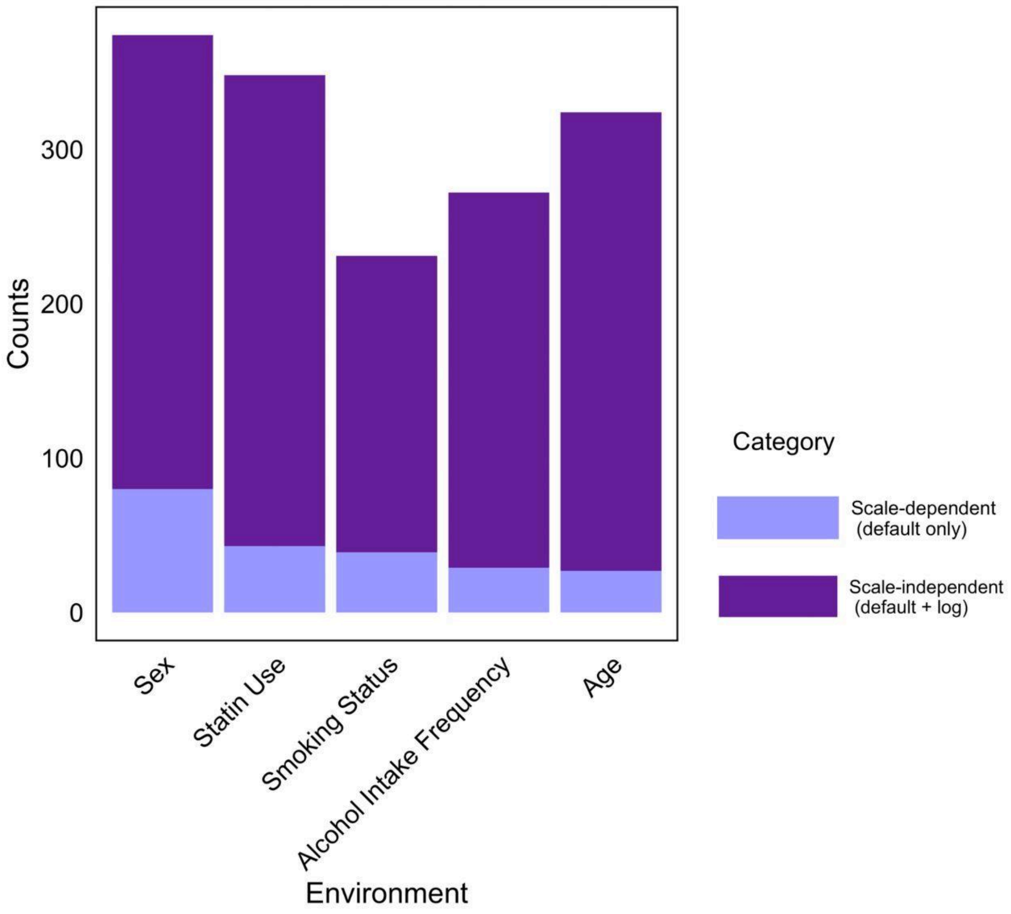


Using a threshold of p=0.05 to determine which interactions are scale-dependent and scale-independent, we find that for sex there are 374 significant interactions (29.7% of total tests). 21.4% of these are eliminated by the log scale, relative to 23.4% when using Bonferroni correction.

### Supplementary Figure 12: Coefficient of Variation and Heteroskedasticity

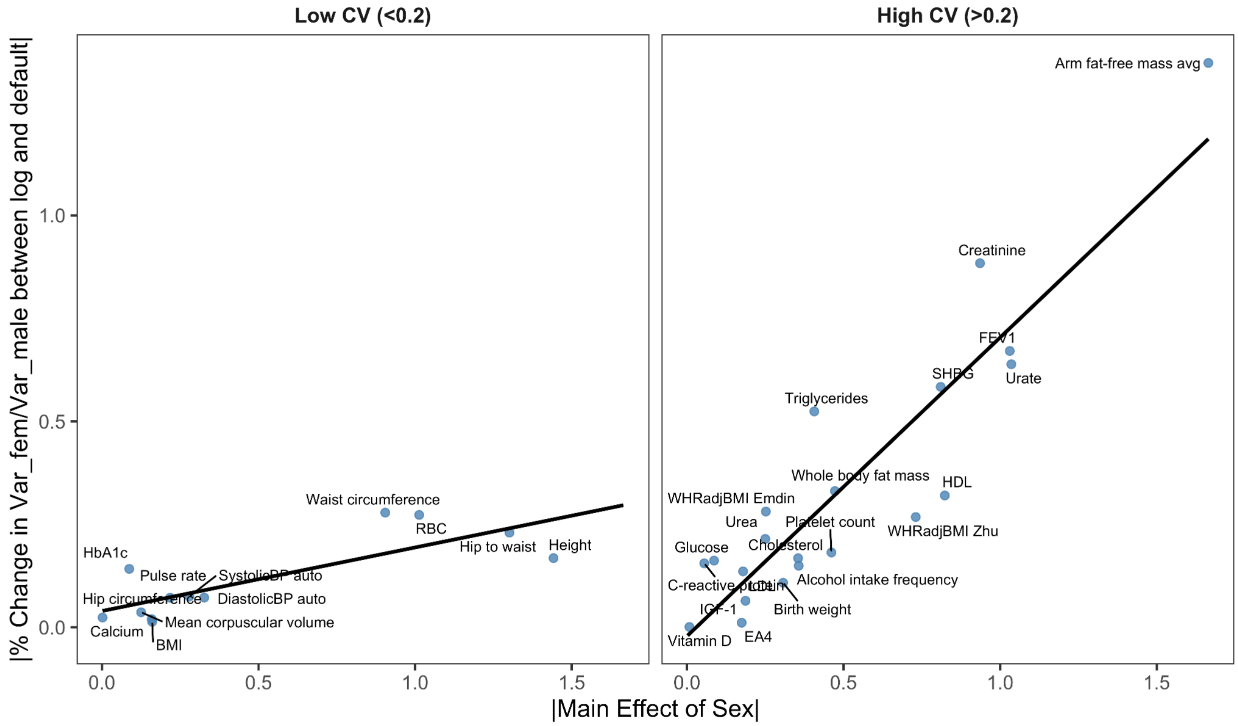


<u>Figure S12.A:</u> There is a positive relationship between the main effect of sex and the change in heteroskedasticity between the two scales, but only when the coefficient of variation is high (>0.2), as was determined by Falconer.

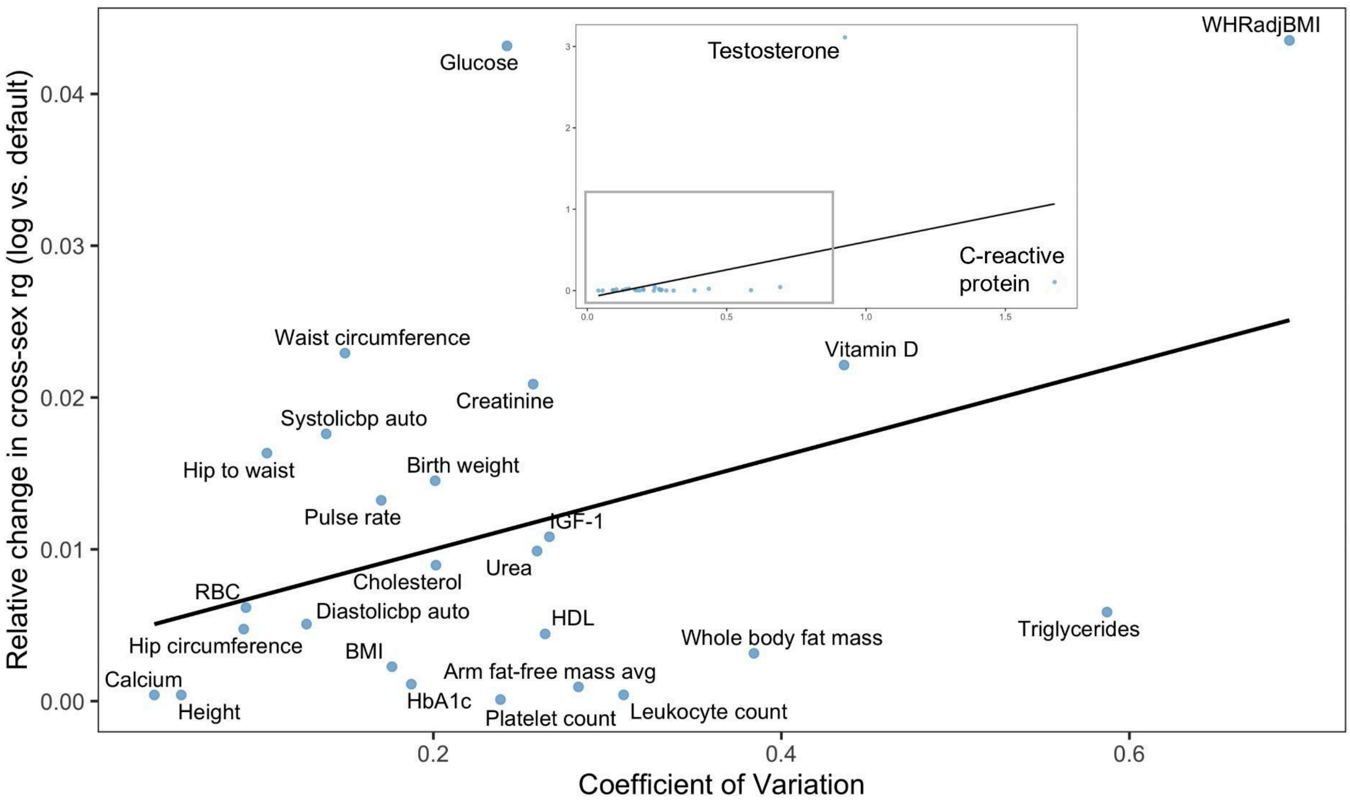


Figure S12.B: Significant relationships between the coefficient of variation and the relative change in cross-sex genetic correlation from default scale to the log scale (p=0.045). Since the Y axis is a ratio, phenotypes with a small $R^2$ on the default scale are positive outliers.

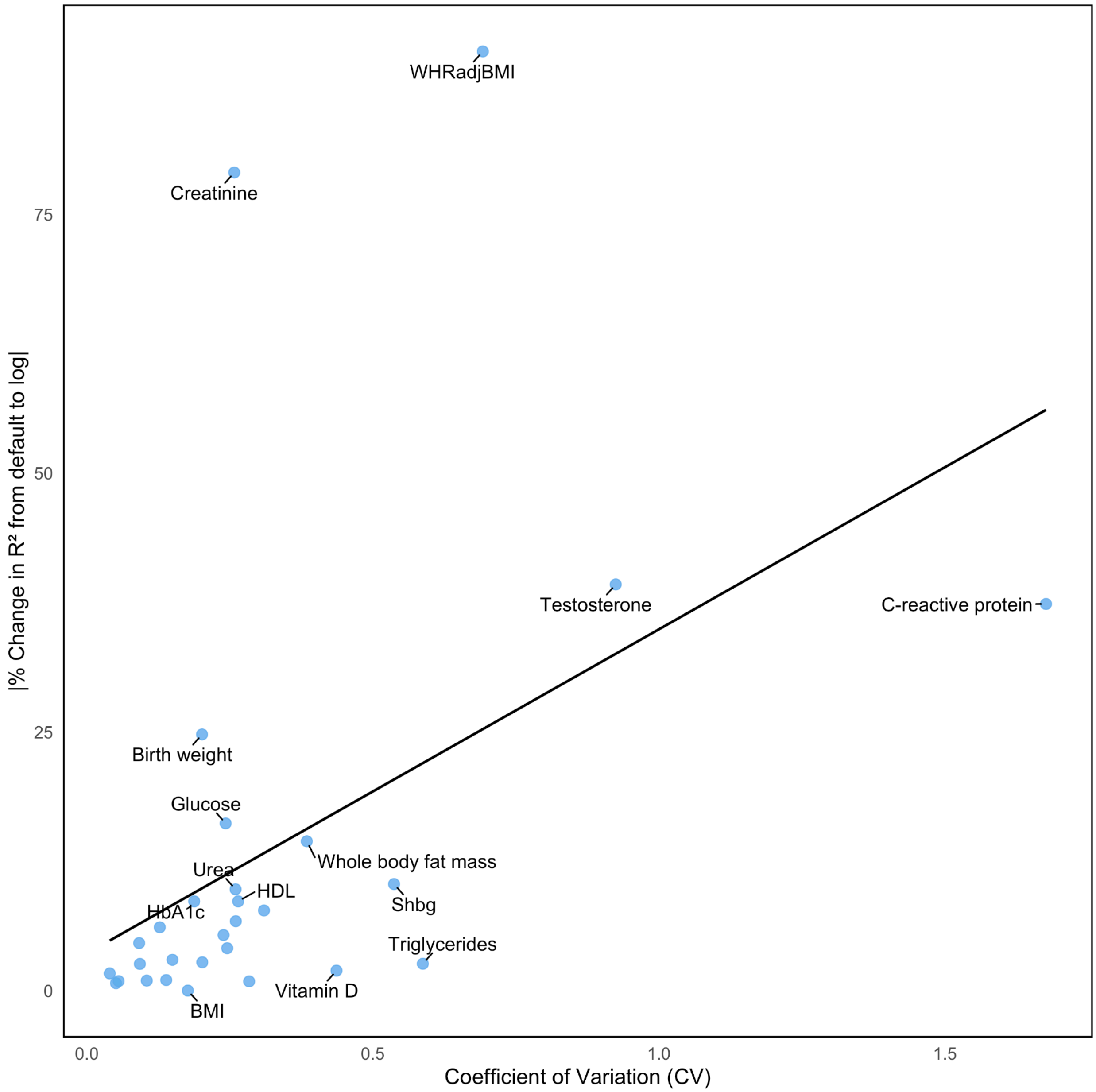


Figure S12.C: Significant correlation between the coefficient of variation and the absolute value of the change in prediction accuracy from the default scale to the log scale (p=1.08e-0.2). We measured prediction accuracy using Pearson $R^2$.

### Supplementary Figure 13: Binary traits simulation

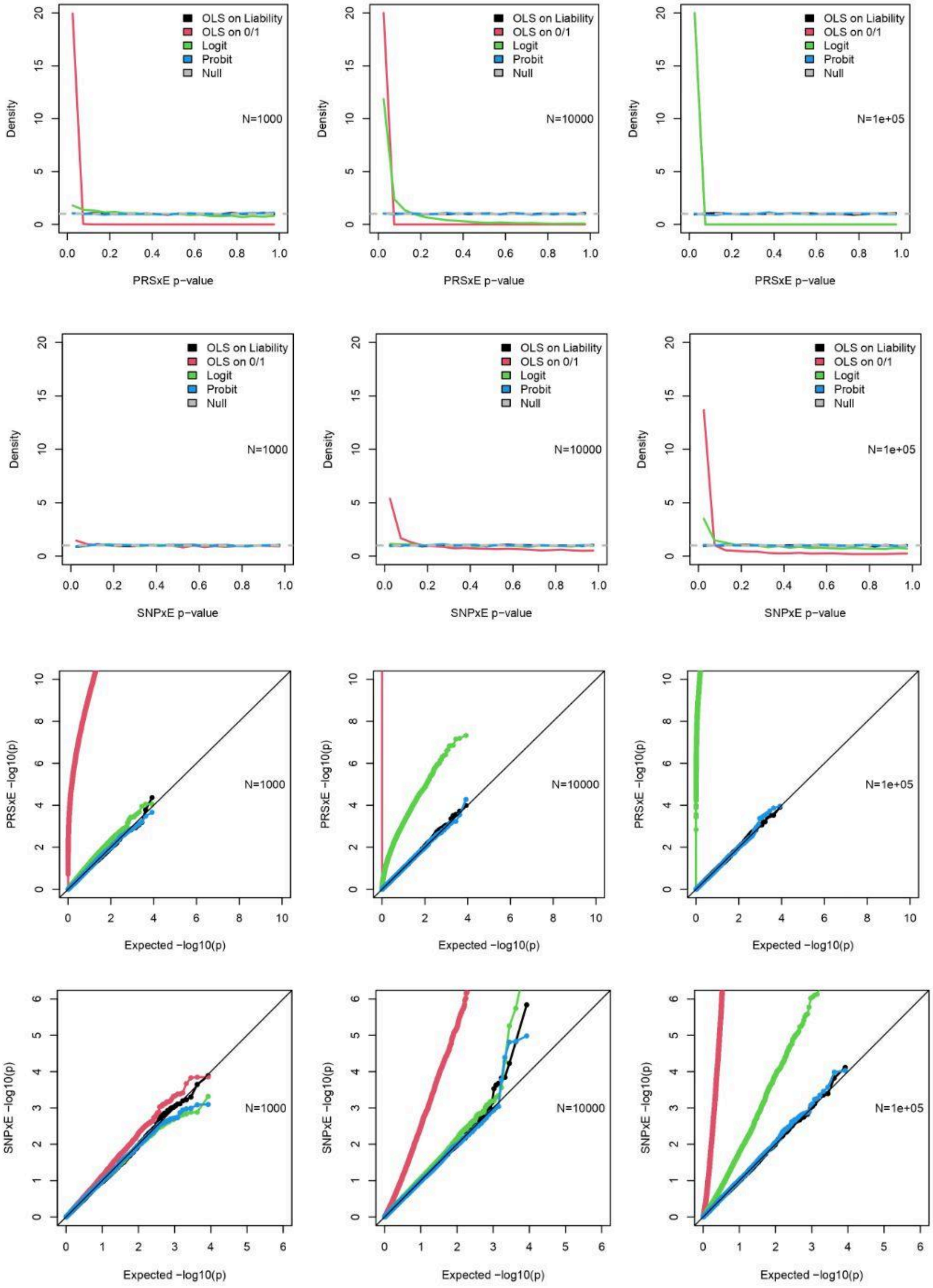

Figure S13: We simulate the liabilities of additive binary traits using either one PGS (top two rows) or 100 SNPs (bottom two rows). The heritability is set at 0.2, the variance explained by the environment is 0.2, the prevalence of the trait is 0.3 and the sample size varies from 1000 to 100,000. We examine the p-value of either the PGS×E or one SNP×E interaction term using four different models: linear regression on the continuous liabilities, linear regression on the binarized phenotype, GLM with a logit link and GLM with a probit link. With large enough samples, we find that GLM with a logit link and linear regression on the binarized phenotype give inflated p-values in both simulation setups.

**Supplementary Figure 13: Main Effect of Each Environment on Each Phenotype**

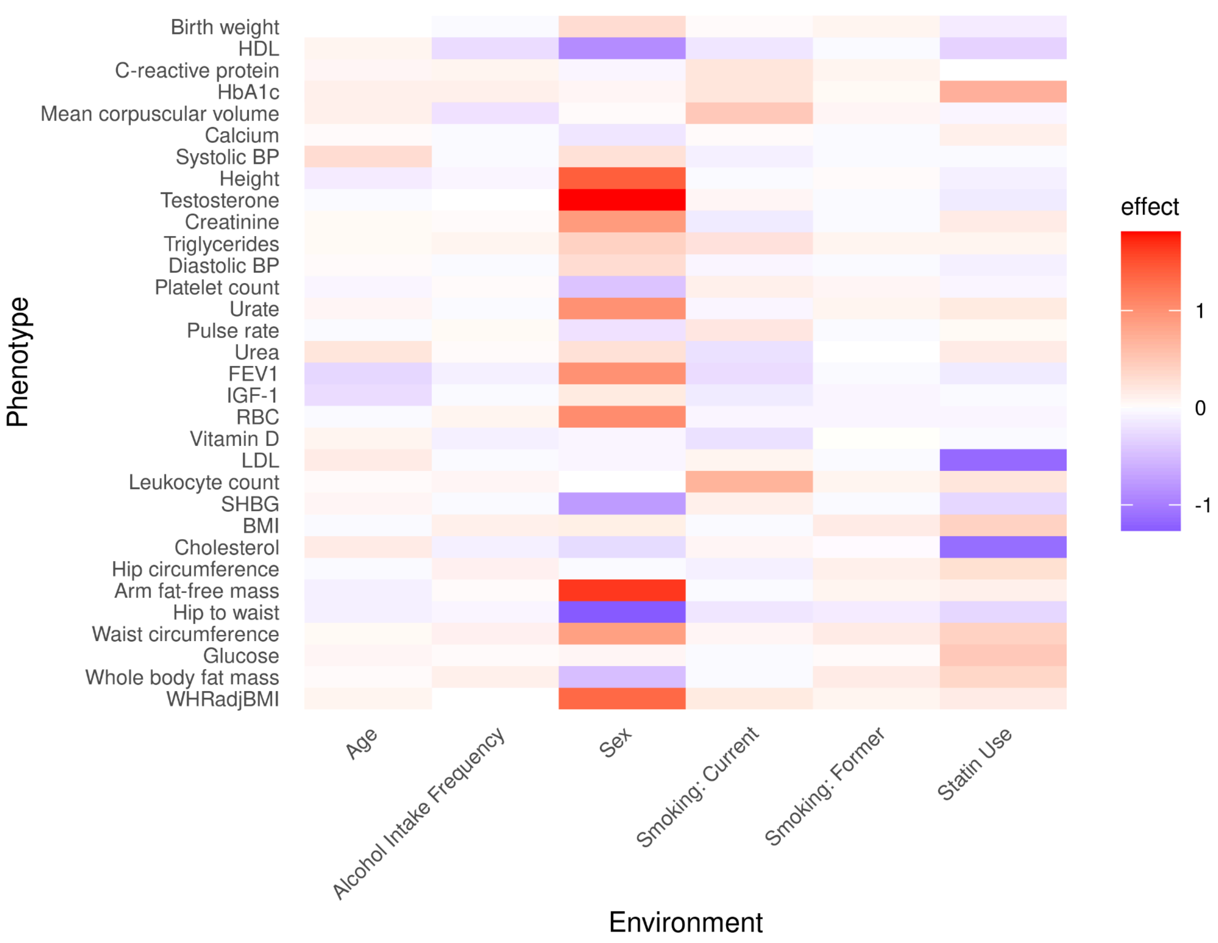

**Supplementary table 1: Environments**

| Environment | Type | Field ID(s) |
|---|---|---|
| Sex | Binary | 31 |
| Age | Continuous | 21003 |
| Statins | Binary | 20003 |
| Smoking status | Categorical | 20116 |
| Alcohol intake frequency | Continuous | 1558 |
| BMI | Continuous | 21001 |

**Supplementary table 2: Phenotypes**

| Phenotype | Name in UK Biobank | Field ID(s) |
|---|---|---|
| Arm fat-free mass avg | Arm fat-free mass (left), Arm fat-free mass (right) | 23125, 23121 |
| Birth weight | Birth weight | 20022 |
| BMI | Body mass index (BMI) | 21001 |
| C-reactive protein | C-reactive protein | 30710 |
| Calcium | Calcium | 30680 |
| Cholesterol | Cholesterol | 30690 |
| Creatinine | Creatinine | 30700 |
| Diastolic BP | Diastolic blood pressure, automated reading | 4079 |
| FEV1 | Forced expiratory volume in 1-second (FEV1) | 3063 |
| Glucose | Glucose | 30740 |
| HbA1c | Glycated haemoglobin (HbA1c) | 30750 |
| HDL | HDL cholesterol | 30760 |
| Height | Standing height | 50 |
| Hip circumference | Hip circumference | 49 |
| Hip to waist | Hip circumference, Waist circumference | 49, 48 |
| IGF-1 | IGF-1 | 30770 |
| LDL | LDL direct | 30780 |
| Leukocyte count | White blood cell (leukocyte) count | 30000 |
| Mean corpuscular volume | Mean corpuscular volume | 30040 |
| Platelet count | Platelet count | 30080 |
| Pulse rate | Pulse rate | 4194 |
| RBC | Red blood cell (erythrocyte) count | 30010 |
| SHBG | SHBG | 30830 |

| Phenotype | Name in UK Biobank | Field(s) |
|---|---|---|
| Systolic BP | Systolic blood pressure, automated reading | 4080 |
| Testosterone | Testosterone | 30850 |
| Triglycerides | Triglycerides | 30870 |
| Urate | Urate | 30880 |
| Urea | Urea | 30670 |
| Vitamin D | Vitamin D | 30890 |
| Waist circumference | Waist circumference | 48 |
| Whole body fat mass | Whole body fat mass | 23100 |
| WHRadjBMI | Waist circumference, Hip circumference, Body mass index (BMI) | 48, 49, 21001 |

**Supplementary table 3: Polygenic Scores**

| Polygenic Score | Name in UK Biobank (Standard PRS for …) | Field(s) |
|---|---|---|
| Age menopause | Age at menopause (AMA) | 26202 |
| Alzheimer's | Alzheimer's disease | 26206 |
| Arthritis | Rheumatoid arthritis (RA) | 26273 |
| Atrial fibrillation | Atrial fibrillation (AF) | 26212 |
| Bipolar | Bipolar disorder (BD) | 26214 |
| BMI | Body mass index (BMI) | 26216 |
| Bone mineral density | Estimated bone mineral density t-score (EBMDT) | 26234 |
| Bowel cancer | Bowel cancer (CRC) | 26218 |
| Breast cancer | Breast cancer (BC) | 26220 |
| Crohn's | Crohn's disease (CD) | 26229 |
| Coeliac | Coeliac disease (CED) | 26225 |
| Coronary artery | Coronary artery disease (CAD) | 26227 |
| CVD | Cardiovascular disease (CVD) | 26223 |
| Glaucoma | Primary open angle glaucoma (POAG) | 26265 |
| HBA1C | Glycated haemoglobin (HBA1C_DF) | 26238 |
| HDL | High density lipoprotein cholesterol (HDL) | 26242 |
| Height | Height (HEIGHT) | 26240 |
| Hypertension | Hypertension (HT) | 26244 |
| Intraocular pressure | Intraocular pressure (IO) | 26246 |
| LDL | Low density lipoprotein cholesterol (LDL) | 26250 |
| Lupus | Systemic lupus erythematosus (SLE) | 26278 |
| Macular degeneration | Age-related macular degeneration (AMD) | 26204 |
| Melanoma | Melanoma (MEL) | 26252 |
| MS | Multiple sclerosis (MS) | 26254 |

| **Polygenic Score** | **Name in UK Biobank (Standard PRS for …)** | **Field(s)** |
|---|---|---|
| Osteoporosis | Osteoporosis (OP) | 26258 |
| Ovarian cancer | Epithelial ovarian cancer (EOC) | 26232 |
| Parkinson’s | Parkinson’s disease (PD) | 26260 |
| Prostate cancer | Prostate cancer (PC) | 26267 |
| Psoriasis | Psoriasis (PSO) | 26269 |
| Schizophrenia | Schizophrenia (SCZ) | 26275 |
| Stroke | Ischaemic stroke (ISS) | 26248 |
| T1D | Type 1 diabetes (T1D) | 26283 |
| T2D | Type 2 diabetes (T2D) | 26285 |
| Ulcerative colitis | Ulcerative colitis (UC) | 26287 |
| Venous thromboembolic | Venous thromboembolic disease (VTE) | 26289 |
